# Structure and signaling mechanism of *Helicobacter pylori* transducer-like protein D

**DOI:** 10.64898/2026.01.16.699579

**Authors:** Kailie Franco, Megan Dilorio, Adam J. Simpkin, Ronan M. Keegan, Karen Kallio, Emily Goers Sweeney, Michelle Colbert, S. James Remington, Arkadiusz W. Kulczyk, Arden Baylink

## Abstract

Here, we present the first high-resolution structures of an intact extended coiled-coil signaling system, the chemoreceptor transducer-like protein D (TlpD) from *Helicobacter pylori*, determined by X-ray crystallography in two crystal forms at 2.4-3.3 Å resolution and cryo-EM at 4.2 Å resolution. The structures capture multiple signaling conformations and reveal how small perturbations within the ligand-binding domain are propagated through the extended coiled-coil to regulate the kinase interface more than 140 Å away. Signal transmission is mediated by a cascade of side-chain rearrangements that coordinate gear-like axial rotation of the interlocked α-helices and remodel conformation and order of the distal hairpin tips. These near-atomic resolution structures provide a framework for understanding the mechanisms of long-range allostery and cooperativity in chemoreceptors and, more broadly, information transfer in coiled-coil sensory systems.

## Introduction

Coiled-coil protein assemblies are elongated, intertwined α-helical structures that comprise approximately 5% of all translated proteins and serve as ubiquitous scaffolds for cellular signaling processes ^1–3^. Cells employ these sensory systems to perceive physicochemical stimuli in their environment and control adaptive responses in motility, metabolism, gene regulation, and division ^4–7^. Coiled-coils are characterized by repeating amino acid motifs, most commonly with heptad periodicity and standard positions denoted as *a-g* ^8,9^. Mutations or defects in coiled-coil functions are implicated in a range of diseases, including muscular dystrophy, cardiomyopathy, neurodegenerative disorders, and cancer ^5,6,10–13^. Canonically, coiled-coil signaling systems comprise a ligand-binding domain (LBD) that controls the conformation and dynamics of the extended coiled-coil architecture to transduce signals over distances spanning >100 Å and regulate distal functional outputs ^4,14,15^. It has been proposed that such signals may be relayed through coupled entropic switches, in which a stimulus redistributes conformational order and disorder (entropy) among oppositely coupled segments of the coiled-coil, rather than a toggling between two rigid conformations ^16^. Absence of high-resolution structural information has hampered understanding of the structural mechanisms by which coiled-coil sensory proteins integrate and prioritize signaling inputs and administer long-range allosteric control, as many are transmembrane and recalcitrant to crystallization and other structure determination methods in their full-length forms.

To circumvent the long-standing challenges associated with structural studies of transmembrane coiled-coil signaling systems, we focused on a conserved class of cytoplasmic, or “soluble” chemoreceptors that retain core signaling architecture but lack membrane-spanning segments ^17–19^. Chemoreceptors are α-helical, antiparallel coiled-coil sensors that detect nutrients and other effectors, allowing bacteria to spatially interpret their environment and direct swimming motility through chemotaxis ^4,17,20–25^. *In vivo* they assemble into trimers-of-homodimers and higher-order signaling networks known as nanoarrays, but ligand sensing and signal transduction are encoded within the receptor homodimer itself, making it a tractable unit for mechanistic study ^26^. The sensitivity and selectivity of chemoreceptors for many different types of biological stimuli has also motivated development of synthetic reporter systems and chemotaxis-based tools for microbe-delivered therapeutics ^27,28^, and they represent potential targets for antimicrobial intervention and microbiome engineering ^20,22^.

We selected for study the cytoplasmic chemoreceptor TlpD from *H. pylori*, which is regulated by a C-terminal CZB domain and plays important roles in gastric colonization and infection ^29–35^. The CZB domain is a type of LBD found in chemoreceptors and diguanylate cyclases that binds a single Zn² ion through three non-contiguous histidines and a cysteine thiolate, mediating behavioral responses to stimuli that alter zinc homeostasis ^29,30,34–39^. For some host-associated bacteria, CZBs function as sensors of host inflammatory processes, and the Cys-Zn moiety can be selectively oxidized by the neutrophil-derived oxidant HOCl to form cysteine-sulfenate (Cys-SOH), promoting Zn² release and regulating chemotaxis and biofilm formation ^33–35,38^.

We report herein the first structures of TlpD, the longest full-length extended coiled-coil sensory system solved at high resolution to date. Two crystal structures of TlpD from *H. pylori* strain J99 were solved: a *P*1 structure containing two homodimers at 2.45-3.28 Å resolution, and a *P*2_1_2_1_2_1_ structure containing one homodimer at 2.92-3.42 Å (Fig. 1a-c, Extended Data Fig. 1-7, Supplemental Table 1, Supplemental Movie 1, Methods). In parallel, we applied cryo-EM to solve a 4.2 Å structure of TlpD from strain SS1, the highest resolved cryo-EM structure for a chemoreceptor thus far (Fig. 1d-f, Extended Data Fig. 7-8, Supplemental Table 2, Supplemental Movie 1, Methods). Together, the four structures represent plausible signaling intermediates and provide high-resolution snapshots of long-range signal transduction of a coiled-coil signaling assembly (Fig. 1).

**Fig. 1.**
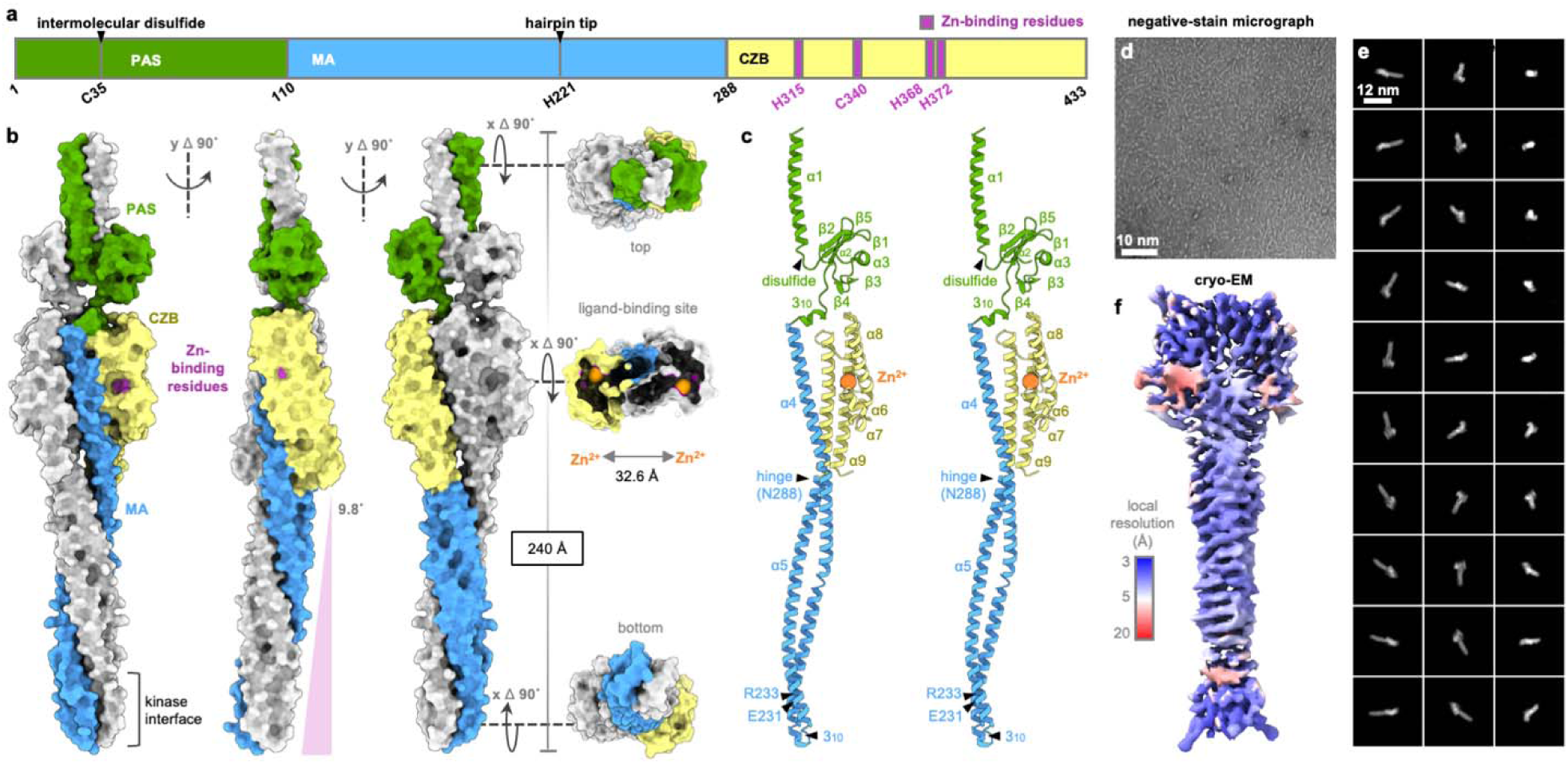
Architecture of TlpD. **a**. Domain organization of TlpD, with key structural features indicated. **b**. Crystal structure of the TlpD homodimer in six orientations (*P*1 chains CD), with one protomer colored as in a and the partner chain in white. The molecular surface is shown, with Zn²□ ions depicted as enlarged orange spheres. **c**. Cross-eyed stereo view of a TlpD monomer, highlighting secondary structure elements and key structural features. **d.** Representative negative-stain micrograph showing approximately 500 TlpD particles dispersed across the field of view. **e.** Representative cryo-EM 2D class averages of the TlpD homodimer. **f.** Local resolutions mapped onto the Coulomb map surface.

## Results

### Architecture of the TlpD homodimer

The TlpD homodimer is revealed to form an elongated key-like structure, 240 Å in length, in which the three domains of each protomer—an N-terminal Per-Arnt-Sim (PAS) (residues 1-110), methyl-accepting (MA) (111-288) that contains the coiled-coil, and C-terminal CZB (289-433)—are intertwined along the central coiled-coil in a nearly two-fold symmetrical embrace (Fig. 1a-b, Extended Data Fig. 6, Supplemental Movie 1). The protein contains nine α-helices, two 3_10_-helices, and five β-strands, although the secondary structure varies slightly across the chains (Fig. 1c, Extended Data Fig. 3-4). The major structural difference across the TlpD dimers is the degree of bending of the MA domains, ranging from 1.1-11° deviation from the central axis, and is enabled by a structural hinge near residue N288 (Fig. 1c, Fig. 2a, Extended Data Fig. 2c, 5-6, Supplemental Movie 1). The conformational heterogeneity in the MA domains among the TlpD dimers is obvious through superposition of each chain with the dimers aligning with a RMSD of 2.85 Å across 375 Cα positions (Fig. 2a, Extended Data Fig. 9, Supplemental Table 3). The dimer interface is distributed along the length of the protomers and buries a substantial amount of surface area, with 1790 Å^2^ contributed from the N-terminal region, 3310 Å^2^ from the MA domain, and 650 Å^2^ from the CZB domain, and exhibits the prototypical knobs-into-holes packing and hydrophobic seam of coiled-coil oligomers (Fig. 2b, Extended Data Fig. 6) ^1,2,8,9^.

**Fig. 2.**
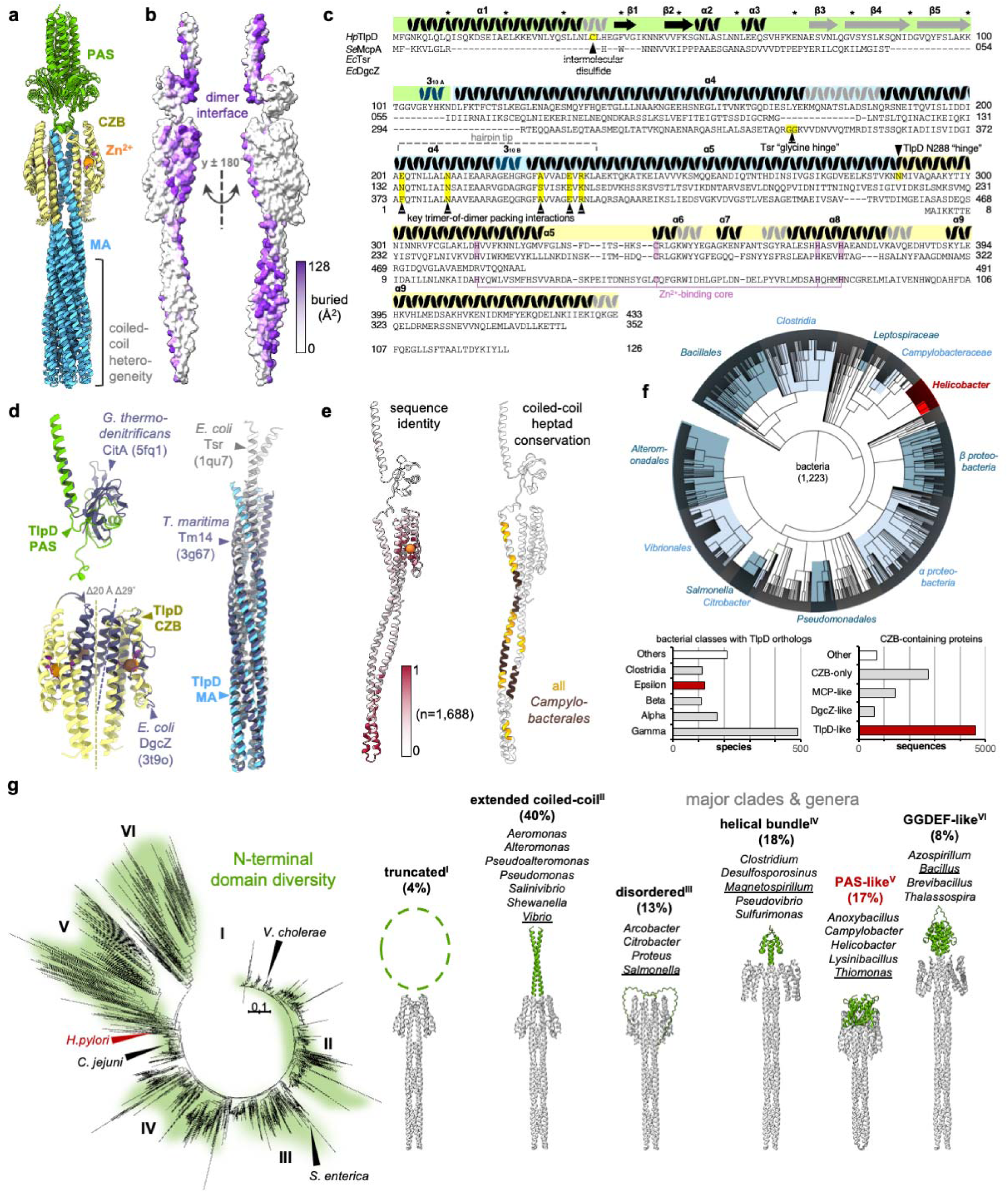
Structural comparisons and evolutionary conservation of TlpD. **a.** Overlay of six TlpD crystal structure chains using residues 115-135, the central core of the MA domain that threads between the CZBs. **b.** The TlpD homodimer interface shown from the *P*1 CD homodimer. **c.** Sequence alignment of *H. pylori* TlpD (strain J99) and related proteins *S.* Typhimurium McpA, another CZB-regulated chemoreceptor, the transmembrane chemoreceptor *E. coli* Tsr, and *E. coli* DgcZ. Helical and strand regions in TlpD are indicated with coils and arrows, respectively, with gray indicating variable regions. Domain regions are colored as in Fig. 1 panel a. Pink indicates the zinc-binding residues, and key regions of structural interest are denoted with yellow highlights and black carrots. **d.** Overlays of relevant structures with TlpD domains, as indicated. **e.** Amino acid conservation across TlpD orthologs mapped onto the monomeric TlpD structure (left). Heptad conservation across all TlpD-like proteins (gold) and across the order *Campylobacterales* (brown) mapped onto the monomeric TlpD structure (right). **f.** Distribution of TlpD orthologs in nature. Top; phylogenetic tree of TlpD-containing organisms, bottom; major TlpD-containing clades and architecture type of CZB-containing proteins. The taxa/architecture groups to which *H. pylori* TlpD belongs are noted in red. **g.** LDDT-based phylogenetic tree based on structures generated from Alphafold 3 of all TlpD-like cytoplasmic CZB-containing proteins using *P*1 chain C as a template (left). Groups are classified based on N-terminal structure, with the percentage of total proteins and representative genera for each group indicated; the genera corresponding to the representative structure are underlined (right).

### Domain structure of TlpD

Sequence alignments of TlpD with three comparators provide context for shared and dissimilar features relative to proteins with homologous domains: (1) the chemoreceptor McpA from *S.* Typhimurium, a CZB-regulated chemoreceptor with similar architecture to TlpD ^39^, (2) the more distantly related *E. coli* chemoreceptor Tsr, which lacks a CZB and is membrane-bound ^40,41^, and (3) diguanylate cyclase Z (DgcZ) from *E. coli*, which has an N-terminal CZB and lacks a coiled-coil (Fig. 2c-d) ^35,36^.

That TlpD has an N-terminal PAS was unexpected based on sequence alone (Fig. 1a-c, Extended Data Fig. 3). The PAS makes little contact with the rest of the protein, joins to the MA domain through a 3_10_-helix, buttresses the α8-α9 loop of the CZB, and is poorly ordered as evidenced by high B-factors and weak electron density in the crystal structures (Extended Data Fig. 2,3,5, Supplemental Movie 1); it was unresolved in the cryo-EM structure (Fig. 1b-c, Extended Data Fig. 7,8). The PAS lacks a definitive ligand-binding pocket, suggesting its function could be mechanical rather than sensory, with its closest analogue being a *Geobacter* histidine kinase (PDB 5fq1; 16% sequence identity; DALI Z-score 6.9; Fig. 1b-c, Fig. 2d). The α1 helix contains an intermolecular disulfide bond at C35, which explains covalently-linked TlpD oligomers seen during our protein purification and in earlier work (Extended Data Fig. 3b, Supplemental Movie 1) ^19,29,30,34^. Whereas Cys residues outside of the LBD are rare in other chemoreceptors (13% of Tsr orthologs), they are present in most TlpD receptors (86%), and might be involved in redox-based regulation and/or provide means for sensory adaptation (Supplemental Data 1) ^40^.

The central MA domain of TlpD adopts the archetypal antiparallel coiled-coil common to the chemoreceptor protein family, whereby the distal coiled-coil hairpin tips serve as the regulatory interface with the histidine kinase CheA (Fig. 1b-c, Fig. 2d, Extended Data Fig. 3, Supplemental Movie 1) ^4,14^. The N-terminal α4 winds down to its hairpin tip and 3_10_-helix, with its apex at H221, then folds back upon itself and ascends through α5 to join to the C-terminal CZB domain (Fig. 1a-c, Fig. 2d). The TlpD MA contains approximately 20 heptads, with stutters appearing at positions 221 and 226, a stammer at 217, skips at 123, 135, 158, 170, 184, 235, 249, 263 and 270 and non-canonical residues appearing in core (*a/d*) positions at 155, 229, 243 and 257 (Fig. 2e). Differences in the MA bending and supercoiling among the TlpD homodimers facilitate large alterations in the positioning of the hairpin tips, although the local structure is retained across residues N205-K234, which twist as a singular module (Fig. 2a, Extended Data Fig. 9-10). Structural heterogeneity in this region is further reflected by the gradient of local resolutions across the cryo-EM maps (Fig. 1f, Extended Data Fig. 7-8). Comparing the structure of the TlpD MA to crystal structures from the cytosolic fragment of Tsr of *E. coli*, and cytosolic receptor Tm14 of *Thermotoga maritima*, which lacks a LBD, we find all three proteins share a conserved topology, especially at the hairpin tips, overlaying within 1.32 Å RMSD over 85 Cα ^41,42^, and TlpD possesses the conserved residues E231 and R233 that mediate the formation of the trimers-of-homodimers signaling unit (Fig. 1c, Fig. 2d, Extended Data Fig. 9s, Supplemental Table 3, Supplemental Movie 1) ^15,41,43^. A notable difference from transmembrane chemoreceptors is that TlpD lacks the six canonical methylation sites used for sensory adaptation in some bacteria, consistent with the absence of CheR and CheB in *H. pylori* (Fig. 2c).

Lastly, the C-terminal CZB domain adopts a compact antiparallel helix bundle formed from α5-α9 (Fig. 1b-c, 2d, Extended Data Fig. 3, 4c-i). TlpD represents the most common CZB protein domain architecture but how the CZB domains of chemoreceptors in their native homodimeric arrangement accommodate the MA coiled-coil was unknown (Fig. 2d) ^34–36^. The structures reveal that this is accomplished by shifting the individual CZBs 9 Å apart so that they straddle the central coiled-coil, which passes between them, while the two-fold symmetry axis at the interface is tilted by approximately 30° relative to DgcZ (Fig. 2d, Extended Data Fig. 9k-l).

### Conservation and phylogenetic distribution of TlpD orthologs

We used phylogenetic and sequence conservation analyses to define the evolutionary constraints imposed by the cooperative functioning of the three domains together as an integrated coiled-coil signaling assembly ^44^. The biological distribution of TlpD spans 1,223 bacterial species and diverse taxa, particularly common in *Alpha*-, *Beta*-, *Gamma*-, and *Epsilonproteobacteria*, as well as in *Clostridia*, *Bacillales*, and *Leptospiraceae* (Fig. 2e-f, Supplemental Data 1). Included among these are many disease-causing bacteria, such as members of the genera *Helicobacter*, *Campylobacter*, *Salmonella*, *Vibrio*, and *Morganella* (Fig. 2f, Extended Data Fig. 9m-r, Supplemental Data 1).

Mapping sequence identity onto the TlpD structure highlights regions preserved across species and broadly essential for function, and reveals striking differences in conservation across the three domains (Fig. 2e). Whereas the CZB zinc-binding core and the MA hairpin tip are strongly conserved at the amino acid level, the extended coiled-coil connecting these signaling input/output regions has undergone extensive evolutionary diversification (Fig. 2e). Despite this sequence variability, key architectural features of the coiled-coil are retained across the protein family, including eight heptad repeats and conserved lesions at positions 135, 170, 229, and 235 (Fig. 2e). These higher-order structural features indicate that the coiled-coil transmission axis acts as a sequence-tolerant scaffold, with signaling properties facilitated primarily by conserved heptads and lesions rather than strict residue-level conservation ^2,45^.

The N-terminal region is important for folding and expression of TlpD, but across homologs is the region of lowest conservation, suggesting it to be a hotspot for evolutionary divergence in this protein family (Fig. 2e) ^46^. As other soluble chemoreceptors utilize their N-terminal regions to interdigitate and form sandwiches of cytosolic signaling arrays, these variations among TlpD orthologs might reflect different modes of array assembly ^47^. To investigate the structural diversity of the TlpD N-terminus, we generated a database of AlphaFold3 homodimeric models for all known orthologs, using our TlpD crystal structure as a template to guide modeling accuracy, and then performed LDDT-based phylogenetic clustering of these models using FoldTree based on N-terminal structure (Fig. 2g, Supplemental Data 1) ^48,49^. This yielded a relatedness tree with six major structural clades. *H. pylori* TlpD is one of the prevalent forms (PAS-like, Clade V, 17%), with the most and least common being the extended coiled-coil (Clade II, 40%) and truncated clades (Clade I, 4%), respectively (Fig. 2g, Supplemental Data 1) ^49^. Clade IV (8%) contains a catalytically inactive GGDEF-like domain and represents an interesting convergence of c-di-GMP signaling and chemotactic regulation within a single chemoreceptor (Fig. 2g).

### Coordination of the Zn^2+^ ligand

Despite being a widespread class of chemosensory domain, CZBs and the structural elements mediating their binding and responsiveness to Zn^2+^ have remained poorly characterized. Fortuitously, the zinc binding site of TlpD is well resolved for all chains of the crystal structures, providing six independent views of the Zn²□ coordination environment (Fig. 3a, Extended Data Fig. 3, 4c-i, Supplemental Movie 1). Chain C of the *P*1 structure provides a particularly exquisite view of the native tetrahedral ligation of the Zn^2+^, with the metal ligand exhibiting a strong 2f_o_-f_c_ electron density peak of approximately 12 σ (Fig. 3a, Supplemental Movie 1). The surrounding helices of the CZB domain cradle the Zn²□ion at the center of the bundle, coordinated by H315 (α5), C340 (α6), and H368 and H372 (α8) (Fig. 3a, Supplemental Movie 1). A salt bridge between D314 and R341 couples the helices α5 and α6 together, and W345 acts as a hydrophobic anchor, securing the α6-α7 segment to the CZB core (Fig. 3a).

**Fig. 3.**
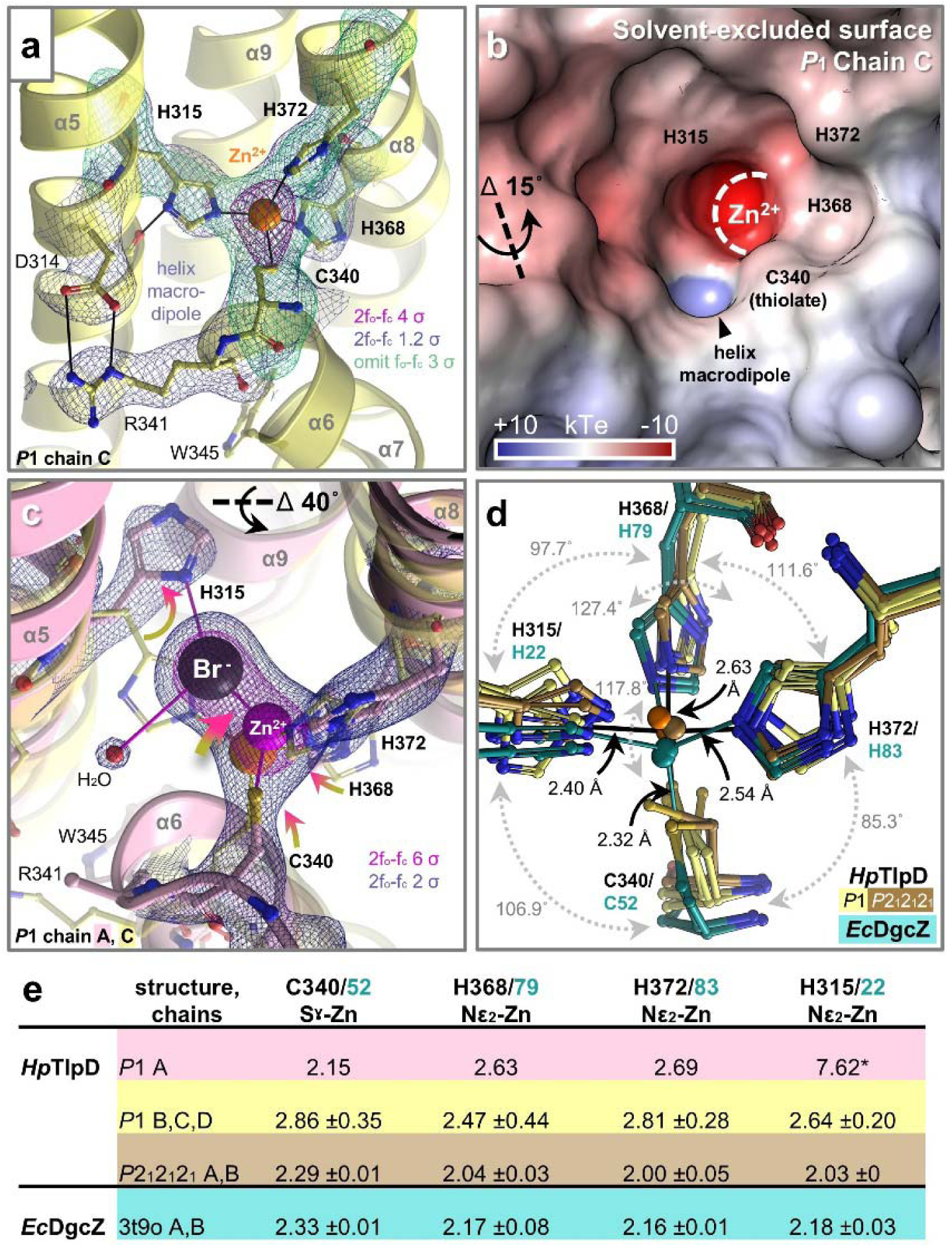
Variable coordination of the Zn^2+^ ligand within the CZB binding site. **a.** Ligand-binding interactions and electron density in *P*1 chain C, as indicated. **b.** Electrostatic surface potential of the zinc-binding site. **c.** Comparison of zinc-binding sites of *P*1 chain C (yellow, “zinc-strained” conformation) with *P*1 chain A (opaque pink, “zinc-disrupted” conformation); structural shifts between chain C and A are indicated by yellow-to-pink arrows. The bromide atom is shown as a black sphere. Electron density mesh is for chain A. **d.** Overlay of zinc-binding sites from the *H. pylori* TlpD *P*1 CD homodimer chains (light yellow), *H. pylori* TlpD *P*2_1_2_1_2_1_ AB homodimer chains (dark yellow, “zinc-intact” conformation), and the two chains of the CZB domain of *E. coli* DgcZ (teal). TlpD residue numbers are indicated in black and DgcZ in teal. Bond lengths and angles shown in black are for *P*1 chain C, and teal bonds for DgcZ chain B. **e.** Zn^2+^ bond lengths ±STD (Å); * denotes distance, not a bond.

TlpD preferentially reacts with HOCl over hydroperoxides via direct oxidation of the zinc-coordinating C340 to form Cys-SOH, promoting release of the Zn^2+^ ligand, and the structures provide new insights into this chemoselectivity ^34,35,38^. A strong helix macrodipole of +10-5 kT/e, generated by the backbone NHs of R341 and L342 at the start of α6, forms a small cavity with strongly positive electrostatic potential near the C340 Sγ (Fig. 3b). This positively-charged pocket could serve as a binding site for HOCl to position the substrate for nucleophilic attack, analogous to the catalytic strategies of thiol peroxidases ^38,50^. Additionally, a halide-binding site is observed within the pocket, with density in the *P*1 dimers consistent with the presence of Cl^-^or Br^-^anions, yielding 2F_o_-F_c_ peaks of approximately 14-18 σ, which could be involved in product release following cysteine oxidation (Fig. 3c, Extended Data Fig. 4c-i).

The signaling state of TlpD is thought to be controlled by Zn^2+^ occupancy, where the ligand-bound form promotes CheA activation (kinase-ON), and the *apo* receptor lowers CheA autophosphorylation (kinase-OFF) ^34,35,39^. Transitional intermediates in this process are modeled by halogen invasion of the binding site, which alters ligand coordination, providing insight into how the overall CZB domain structure is impacted by perturbations in Zn^2+^ ligation (Fig. 3c-d, Extended Data Fig. 4c-i). In some binding sites, halides appear to impose strain on the zinc ligation resulting in unusual coordination geometry, forming a pseudo-trigonal bipyramidal configuration (Fig. 3c-d, Extended Data Fig. 4c-i). The most intrusive occurs in Chain A of the *P*1 structure, where a Br^-^displaces H315, and the lack of the Nε_2_-Zn^2+^ bond is accompanied by a series of structural rearrangements: the H315 sidechain swings toward α9, the Zn²□ shifts by 1.5 Å, and C340, H372, and H368 follow to maintain zinc coordination, while α6 and α8 piston toward the N-terminal domain, altering their register relative to α5 and α9, and these changes in α6 render the R341 sidechain disordered and sever its salt bridge with D314 (Fig. 3c, Supplemental Movie 1). These motions are further supported by the gradient in local resolutions seen in the cryo-EM map, which suggests flexibility in α6, α7, and α8, consistent with these helices forming a structurally dynamic element (Fig. 1f, Extended Data Fig. 7-8, Supplemental Movie 1).

Whereas typical zinc ligation bond lengths for His Nε_2_ and Cys Sγ atoms are 2.0 and 2.3 Å, respectively, the TlpD zinc sites exhibit large variation (Fig. 3c-e) ^51^. Progressively from most distorted to most ideal, the ligation geometries are: (1) the *P*1 Chain A, (2) the *P*1 Chains B, C, D, and (3) the *P*2_1_2_1_2_1_ Chains A and B, which also exhibit the coordination geometry most like that of the DgcZ CZB (Fig. 3d-e, Fig. 5b). These structures suggest that the zinc site forms a metastable coordination network whose plasticity is coupled to the broader conformational state of the CZB domain.

### Ligand-mediated regulation of the MA coiled-coil

The TlpD homodimer structures can be thought of as a continuum of signaling states that we hereafter refer to as the “zinc-intact” (*P*2_1_2_1_2_1_ AB and the cryo-EM structure), “zinc-strained” (*P*1 CD), and “zinc-disrupted” (*P*1 AB) conformations (Extended Data Fig. 2, 4i). These states exhibit varying N-terminal disorder and progressively weaker zinc coordination, accompanied by changes in the structure and dynamics of the MA coiled-coil and hairpin tip regions that regulate signaling output. Together, they provide insight into how sensory inputs from the N-terminal region and CZB zinc-binding site are transmitted through the coiled-coil to modulate receptor output through the structure and relative order of the hairpin tips.

The structural consequences *en route* to ligand release are modeled by the zinc-disrupted conformation (Fig. 3c, Fig. 4a, Supplemental Movie 1). Loss of the H315-Zn interaction is accompanied by an extensive reorganization of 45 residues spanning α6-α7 (Fig. 4a-c). Within this region, W345 (α6) becomes wedged deeper into the helical bundle, bringing it into steric conflict with F307 (α5), which rotates to collide with H135 (α4) (Fig. 4b-c). These changes propagate into deviations in the knob-into-holes packing interactions of α4 that accumulate over a 15-residue span descending toward the hairpin tips, inducing rotation of H151 away from N288′ (α5′) to disrupt an intermolecular hydrogen bond, and providing the basis for the structural hinge that enables bending and flexibility of the MA domain (Fig. 4d-e, Supplemental Movie 1).

**Fig. 4.**
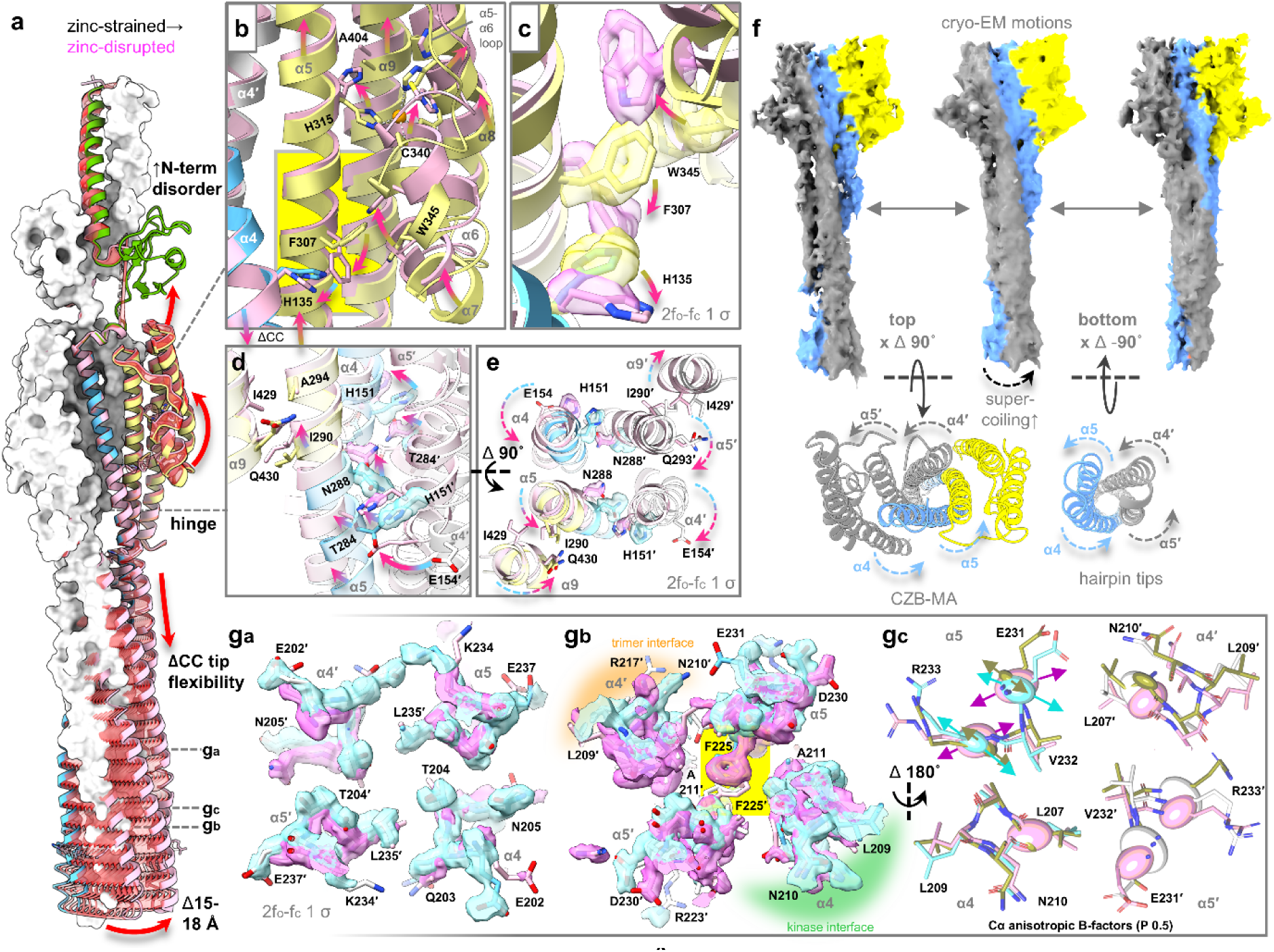
Structural connections between the Zn^2+^-binding site and hairpin tips. **a.** An overlay of the zinc-strained conformation (*P*1 CD, colored by domain as in Fig. 1a and white, respectively) and the zinc-disrupted conformation (*P*1 AB, pink) is shown, with a morph between the two structures highlighting the conformational changes in red. Key structural changes are indicated with red arrows; CC denotes coiled-coil. **b-e.** Close-up views of structural changes in regions of TlpD, as indicated. Arrows highlight shifts from the zinc-strained conformation to the zinc-disrupted conformation; yellow-to-pink arrows in the CZB region and blue-to-pink arrows in the MA regions. **f.** 3D-FLEX and PCA analyses reveal intrinsic dynamics of TlpD homodimers in solution (see also Supplemental Movie 1). **g_a-c_**. Changes in local structure, electron density, and thermal motions in the hairpin tip; electron density is shown as light blue and pink surface for the zinc-strained and zinc-disrupted structures, respectively. Anisotropic B-factors ellipsoids are depicted in g_c_, with ellipsoid probability of 0.5, and arrows highlighting representative differences in thermal motions.

Viewed down the coiled-coil axis toward the hairpin tips, bending at the N288 hinge coincides with asymmetric counterrotation of the coiled-coil helices between the protomers (Fig. 4e, Supplemental Movie 1). These MA-domain rotations are also mechanically coupled to opposing helical twists within the two CZB domains through packing interactions with I290 (α5) and I429 (α9), defining an allosteric regulation network that links the two ligand-binding sites across 30 Å such that conformational changes at one Zn²□ -binding site are coordinated with reciprocal changes at the other (Fig. 4d-e, Supplemental Movie 1). Cryo-EM analyses support the presence of the coupled motions of the CZBs and MA domains in solution, with PCA and 3D-FLEX analyses revealing coordinated CZB rearrangements associated with gear-like rotation of interlocked helices about the central coiled-coil axis (Fig. 4f, Supplemental Movie 1). Spanning 140 Å from sensory input to signaling output, these structural transitions reveal how the localized perturbation at the Zn²□ -binding site is amplified into a receptor-wide signaling response (Extended Data Fig. 10, Supplemental Movie 1).

### Output regulation through conformational entropy

The hairpin tips form the receptor output region within the trimer-of-dimers signaling unit, where their structure and conformational order control kinase activation ^14,15,26,52,53^. Several features of TlpD indicate the hairpin tips occupy a shallow thermodynamic landscape near the transition between alternative states ^54,55^. At the kinase interface, the bundle containing T204 (α4) and L235 (α5) lacks hydrogen-bonding partners capable of satisfying the T204 side-chain hydroxyl, creating a locally unfavorable interaction within the coiled-coil core (Fig. 4g_a_). Similarly, the two-fold symmetry of the TlpD interface positions F225 and F225□ to compete for the same interior packing site, requiring distinct side-chain conformations to satisfy packing interactions of the α5 and α5′ helices (Fig. 4g_b_) ^56^. Based on the orientations observed in the Tsr trimer-of-dimers crystal structure, we infer that the trans/+80° and g□/−30° rotamers denote the protomers forming the kinase and trimerization interfaces, respectively (Fig. 4g_b_) ^56^. Anisotropic B-factors at the hairpin tips exhibit conformation-dependent differences in the magnitude and direction of thermal motions, with the zinc-intact state well ordered, and the zinc-strained and zinc-disrupted states exhibiting larger displacements oriented along different axes (Fig. 4g_c_). Cryo-EM local-resolution analyses further indicate that both the CZB domains and kinase-interface hairpin tip are flexible in solution, supporting a model in which the sensory and output regions of the receptor occupy conformationally labile states poised for signaling transitions (Fig. 1f, Fig. 4f, Extended Data Fig. 7-8, Supplemental Movie 1).

Interestingly, the 2D and 3D analyses of cryo-EM data revealed the presence of higher-order oligomers, including dimers-of-dimers and trimers-of-dimers, even in the absence of CheA and CheW (Fig. 5a, Extended Data Fig. 7-8). To visualize how the TlpD conformational states regulate outputs of the core signaling unit, a trimer-of-dimers model was assembled from each homodimer structure using the Phe225 rotamer assignments to define interior vs. exterior protomer orientations (Fig. 5c) ^41,56,57^. Comparison of the resulting assemblies and crystallographic B-factor patterns provides a rationale for how structural order regulates signal output based on inputs from the N-terminal and CZB regions (Fig. 5c-d). The zinc-intact model exhibits the most ordered hairpin tip region, with the least degree of supercoiling, and the dimers pack well together within the trimer architecture (Fig. 5c-d). Despite similar Zn²□ coordination geometries, the two CZB domains exhibit asymmetric patterns of structural order, with the kinase-interface protomer stabilized and the trimerization-interface protomer destabilized, while the MA hinge regions of both protomers are destabilized (Fig. 5c-d).

**Fig. 5.**
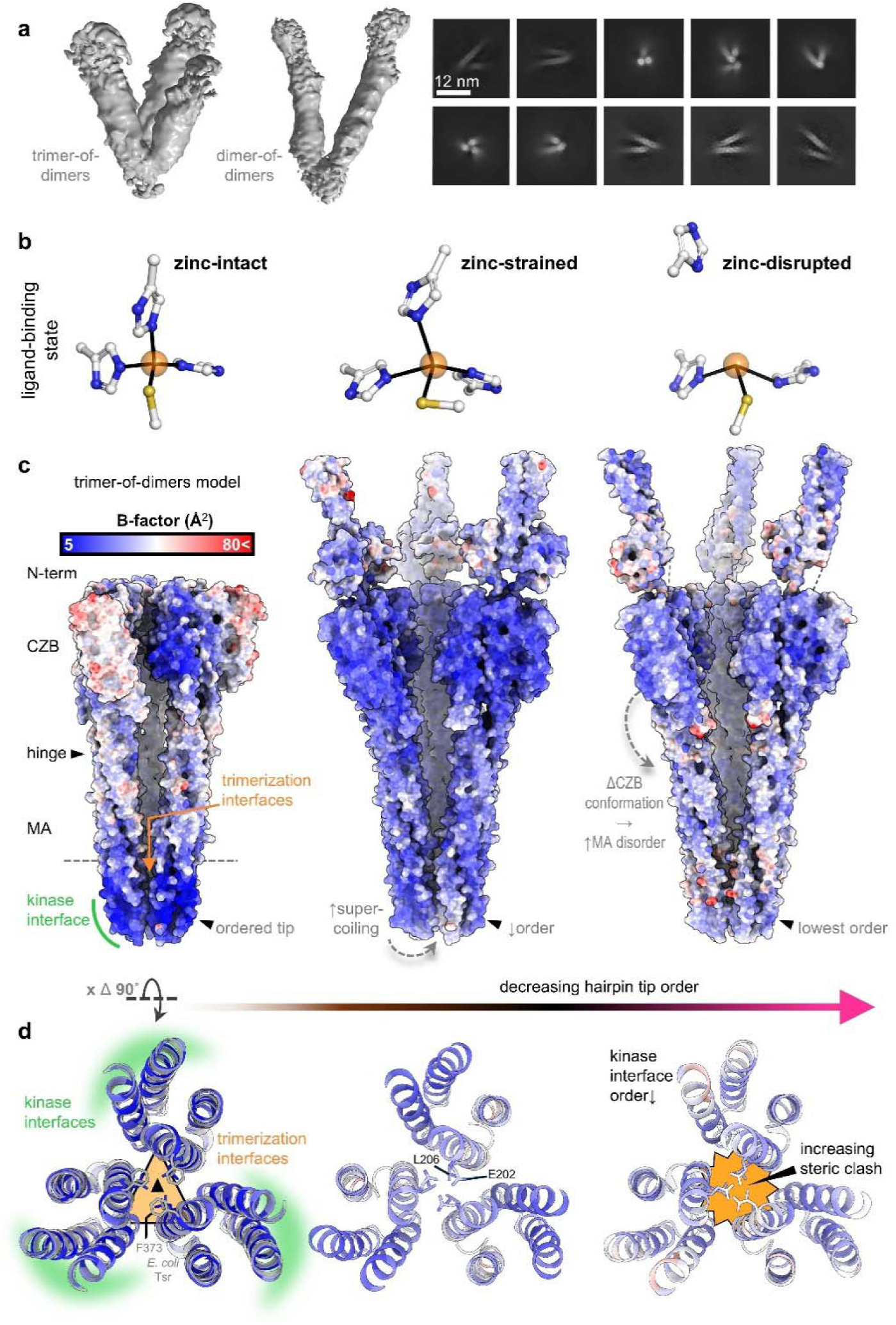
Structural order regulating the trimer and kinase interfaces. **a.** TlpD oligomers observed by cryo-EM. **b.** CZB zinc-binding sites arranged according to increasing perturbation of zinc coordination geometry, representing a progression of signaling states leading toward ligand release. **c.** Models of trimer-of-dimers constructed from each of the TlpD homodimer conformations based on overlays with the *E. coli* Tsr cytosolic fragment (PDB 1qu7) and rendered with surface coloring by B-factor. **d.** Top-down views of the hairpin tip packing, colored by B-factor, with residues involved in potential steric clashes indicated.

The zinc-strained trimer model reveals an additional regulatory role for the N-terminal region. In the only homodimer in which the N-terminal domains are both folded and somewhat ordered, both chains exhibit stabilized CZB domains and MA hinge regions, which may be due to interactions between the PAS and CZB domains (Fig. 5c). Notably, this is the only conformation that captures an intermolecular C35 disulfide bond, suggesting that the N-terminal module may regulate the receptor through redox-dependent disulfide bond formation (Extended Data Fig. 3b) ^30,34^. With this additional regulatory input and stabilized MA hinges, the coiled-coil is more highly supercoiled and the hairpin tips less ordered; in the trimer model, E202 and L206 also have steric clashes at the three-fold interface that could contribute to destabilizing the tips (Fig. 5c–d, Extended Data Fig. 10).

In the zinc-disrupted conformation the hairpin tips are highly supercoiled but the extensive rearrangements in one of the CZB domains, described above, are accompanied by reduced order throughout the receptor and poorly ordered hairpin tips (Fig. 5c-d, Extended Data Fig. 10). In this structure, it appears that the partial release of the Zn^2+^ ligand from one CZB is disruptive to the PAS interface, with a loss of order in the N-terminus of that protomer and corresponding destabilization of the N-terminal region and CZBs of the partner chain (Fig. 5c). The trimer model also predicts substantial steric conflicts to occur at the trimerization interface, further destabilizing the hairpin tips (Fig. 5d). From this series of observations, we infer that the zinc-intact conformation, with the most stabilized hairpin tips, reflects the kinase-ON state, and that the zinc-disrupted conformation, with its destabilized hairpin tips, is shifted toward kinase-OFF, with the zinc-strained conformation representing an intermediate state of activation (Fig. 6a).

**Fig. 6.**
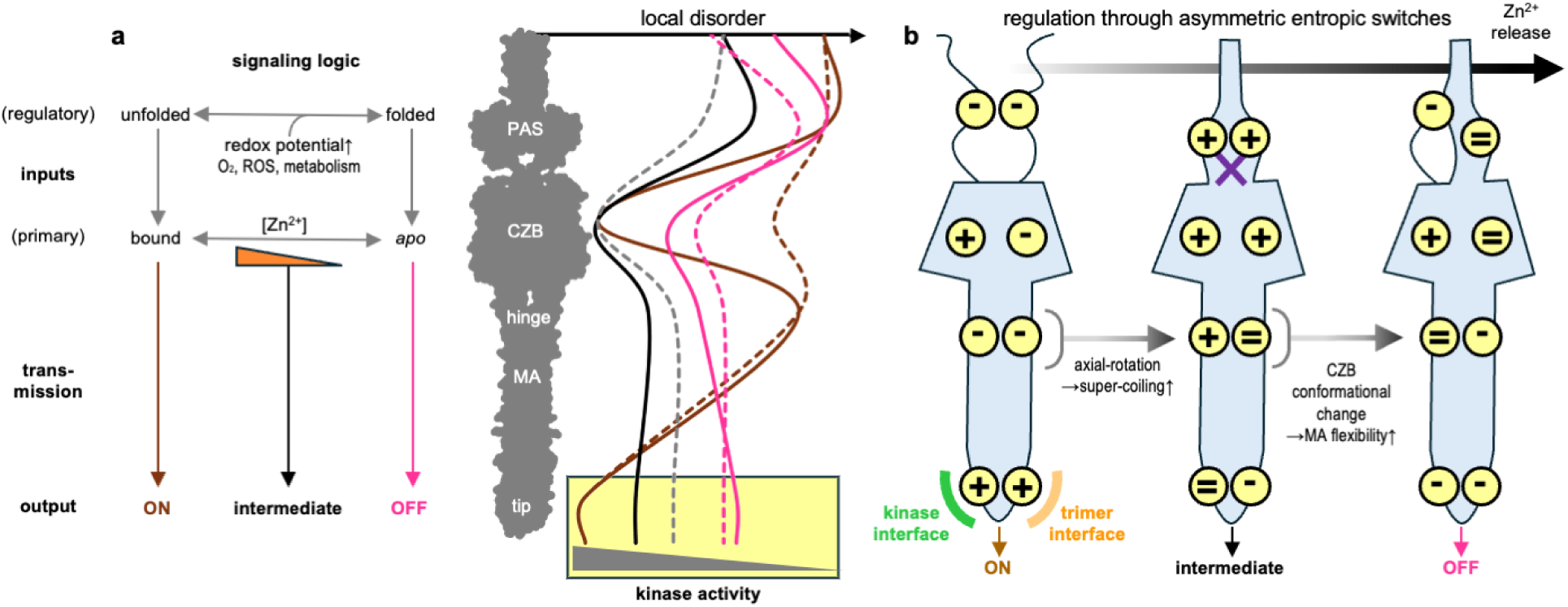
Model for allosteric regulation through asymmetric entropic switches. **a.** Proposed signaling logic of TlpD to generate graded kinase output states. Zn²□ occupancy of the CZB domain serves as the primary regulatory input and is governed by Zn²□ availability, whereas the folded state of the N-terminal PAS domain acts as a secondary regulatory input that is stabilized by intermolecular disulfide formation in response to cellular redox conditions. Signaling output is controlled through shifts in local conformational order that redistribute stability between discrete regions of the receptor and asymmetrically across the homodimer; kinase interface chains shown as solid lines and trimer interface chains as dashed lines in the right panel. **b.** In this model, signal transmission proceeds through asymmetric entropic regulatory switches within the homodimeric receptor, where order is denoted as high (+), intermediate (=), or low (−) for each segment, as indicated. The C35 disulfide is denoted as a purple cross. Progression toward Zn²□ release drives sequential conformational transitions: first, axial rotation of the coiled-coil increases supercoiling, destabilizing the kinase interface and promoting the transition from ON→ intermediate. Release of the zinc ligand triggers conformational changes in the CZB that increases flexibility in the MA hinge, destabilizing the kinase interface. Thus, both oligomeric interfaces of the kinase tip become destabilizing, promoting the transition from intermediate → OFF.

## Discussion

Long α-helical coiled-coils are among the most widespread protein signaling architectures in biology, and mediate information transfer over extended molecular distances ^1–3^. These structures of TlpD provide the first high-resolution views of a full-length long coiled-coil signaling system where specific ligand interactions and individual residue and sidechain motions can be mapped to long-range signal transduction.

TlpD can be thought of as a three-component transponder comprising two input receivers, the N-terminal PAS and C-terminal CZB, and a coiled-coil transmitter (MA domain) that encodes these inputs and transduces the response to the hairpin tips. From our structural studies we derive a two-input signaling model for long-range allostery in this system that operates through asymmetric distribution of order between the protomers of the homodimer (Fig. 6a). The CZB domain functions as the primary sensory module through Zn²□ coordination, with the bound and *apo* states favoring kinase-ON and kinase-OFF signaling, respectively, through regulation of hairpin tip stability (Fig. 6a). This module is highly conserved across orthologs, suggesting that the MA-CZB signaling unit arose early in prokaryotic evolution and has been maintained throughout the diversification of major bacterial lineages (Fig. 2e-f, Fig. 6a, Supplemental Data 1) ^34,35,37,39^. The N-terminal region acts as a secondary layer of regulation whose folded state tempers the dynamics of the hairpin tips by stabilizing the CZB domains and reducing the asymmetric distribution of conformational order across the dimer, as seen from comparison of the zinc-intact versus zinc-strained conformations (Fig. 5, Fig. 6a-b). The presence of the N-terminal intermolecular disulfide bond in the zinc-strained conformation further suggests that in TlpD, this regulatory input is responsive to cellular redox state, and associated physiological cues, such as oxygen availability, reactive oxygen species, or metabolic activity, as also supported by other evidence (Extended Data Fig. 3b) ^34,38,39,50,58^. Hence, the network of allosteric interactions of these two sensory modules is coordinated such that distinct stimuli can be prioritized and integrated into a coherent signaling output (Fig. 6).

The TlpD structures provide a high-resolution system to evaluate the entropic switch model of long-range signal transduction ^26^, and overall our observations reasonably align with this model. For TlpD, perhaps the most critical entropic control point is the reciprocal relationship in conformational order between the MA hinge and hairpin tips (Fig. 5, Fig. 6). However, the experimental structures add an additional dimension by revealing that entropic control can be mediated asymmetrically between the receptor protomers (Fig. 6). This is exemplified by the zinc-intact conformation in which the two CZBs exhibit highly asymmetric order, accompanied by a destabilized MA hinge and ordered hairpin tips (Fig. 5, Fig. 6). Asymmetric behavior within chemoreceptor homodimers has long been implicated in their signaling and negative cooperativity, and these structures provide further direct structural insight into this property by revealing the network of interactions connecting the two ligand-binding sites (Fig. 4d-e) ^56,59–61^. More broadly, our findings support that long-range signaling by extended coiled-coil receptors may be governed as much by the redistribution of conformational entropy as by discrete conformational rearrangements.

## Methods

### Crystallization and structure solution of TlpD

Obtaining well-diffracting TlpD crystals proved exceptionally challenging. We designed numerous constructs and screened thousands of crystallization conditions, yet the protein was uncooperative in forming crystals suitable for high-resolution structure determination (Extended Data Fig. 1). Early trials with full-length TlpD from *H. pylori* strain J99 produced thin, needle-like crystals under many conditions, and occasionally small wedge-shaped crystals with some three-dimensionality, but these could not be reproduced reliably and diffracted poorly (Extended Data Fig. 1a-b). Systematic optimization of protein concentration, precipitants, pH, and temperature, as well as crystal seeding, additive screening, and cryoprotectant treatments, failed to improve crystal growth or diffraction. Parallel attempts with TlpD orthologs from *Salmonella* Typhimurium, *Campylobacter jejuni*, and alternate *H. pylori* strains (SS1 and G27) were unfruitful. After a decade of exhaustive trials exploring standard approaches, high-quality crystals remained elusive.

Remarkably, serendipity provided a breakthrough. Before discarding old crystallization trays set up years earlier, visual inspection revealed a nearly dried hanging drop containing several unusually large, blade-shaped TlpD crystals measuring 0.5-1 mm (Extended Data Fig. 1c). These crystals were extremely fragile and prone to crumbling even with gentle manipulation. The TlpD crystals were soaked with glycerol as a cryoprotectant, and bromine for potential SAD or MAD phasing, before flash freezing for cryogenic X-ray diffraction. Laboratory records revealed the original crystallization condition to be 0.1 M sodium malonate, 0.5 mM TCEP, and 12% PEG 3350, with crystals having grown for approximately two years at 25 °C. All subsequent attempts to reproduce crystals of comparable size and quality in this condition have failed, suggesting these TlpD crystals are the product of exceptionally rare and stochastic circumstances. Likely, this is due to the variation in N-terminal folding, C35 disulfide formation, and differences in Zn^2+^ binding state, which together compound and decrease the likelihood of productive crystal nucleation and ordered growth (Extended Data Fig. 2).

High-resolution diffraction data were obtained from two TlpD crystals, yielding datasets at 2.45-3.30 Å resolution and putative assignments to space groups *P*1 and *P*2□2□2□(Extended Data Fig. 1d-e). Although initially promising, both datasets had substantial anisotropy, and standard approaches for structure determination stalled. Attempts at SAD or MAD phasing failed, and selenomethionine-labeled TlpD did not yield quality crystals sufficient for a derivative. We attempted molecular replacement (MR) using orthologous structures of the MA domain and the single available CZB structure, from *E. coli* diguanylate cyclase Z (DgcZ) ^36,41,42^. Brute-force rotation searches were performed to exploit the low-symmetry of the putative *P*1 lattice, but MR was unsuccessful. Additional strategies, including *ab initio* phasing and MR using models from RosettaMR, RosettaFold, I-TASSER, multiple generations of AlphaFold, and coiled-coil–specific tools such as AMPLE and ARCIMBALDO, were also unsuccessful. As a result, both TlpD datasets remained unsolved for an additional nine years.

The eventual solution came about by employing new prediction tools and a multi-step MR strategy. A homodimeric model of TlpD was generated using ABCFold and Boltz-1 and then segmented with Slice N’ Dice based on regions predicted to be flexible to enable independent MR searches for different portions of the molecule ^64–66^. This approach yielded promising domain placements in the *P*1 dataset (Extended Data Fig. 2). Extensive manual rebuilding ultimately produced a complete solution containing two TlpD homodimers (Extended Data Fig. 2a). This success, in turn, guided solution of the *P*2□2□2□ dataset, which contains a single homodimer in the asymmetric unit (Extended Data Fig. 2b). Retrospectively, the principal obstacle in the MR attempts appears to have been the angular bend in the coiled-coil segment of the MA domain near N288 (Extended Data Fig. 2c); subdividing the coiled-coil into smaller search fragments allowed correct placement of the full assembly.

To address the impact of anisotropy, which contributed to noisy electron density maps and elevated B-factors, we applied the STARANISO pipeline and determined appropriate resolution cutoffs for the *P*1 and *P*2□2□2□ datasets to be 2.45-3.3 Å, with 2.6 Å used for refinement, and 2.92-3.42 Å, and 3.30 Å used for refinement, respectively (Extended Data Fig. 5) ^67^. Although the final R- and B-factors remain higher than typical at these nominal resolutions, the resulting maps are substantially improved, and most regions of the protein are well resolved in all six chains (Extended Data Fig. 3, Supplemental Table 1). Clear electron density is present for the CZB domains, the Zn²□ ligands and coordinating residues, and the kinase-interacting regions (Extended Data Figs. 3, 4c-i). The primary exception is the N-terminal region, which varies among the three homodimers: it is modeled in three of the four chains in the *P*1 structure, while in the *P*2□2□2□ structure crystal packing prevents it from folding, and the region is unresolved (Extended Data Fig. 2-3). For refinement of the zinc-intact structure, strict geometry restraints were employed to overcome the tendency of the model to be distorted by poorly resolved regions within the highly anisotropic data, but, even so, a few geometric outliers remain at residues 143 and 268.

### Negative-stain EM

Samples of TlpD from *H. pylori* strain SS1 were screened by negative-stain EM over a wide range of concentrations obtained by diluting the protein stock (4.8 mg/ml) in 150 mM NaCl, 100 mM Tris pH 7 and 1 mM Tris(2-carboxyethyl)phosphine (TCEP) (Fig. 1d). The 3 µl aliquots were applied to carbon-coated 300-mesh copper grids, which were previously rendered hydrophilic with PELCO easiGlow glow discharger. Grids were stained with 1.5% (w/v) uranyl formate containing 0.125% trehalose for 30 s at room temperature, as described previously ^68–72^. After staining, grids were air-dried and inserted into the microscope. Images were recorded using Philips CM-12 electron microscope operating at 80 kV acceleration voltage at a magnification of 60,000 or 75,000 times corresponding to the pixel size of 1.95 Å/px or 2.44 Å/px, respectively (Fig. 1d).

### Cryo-EM sample preparation, screening and data collection

Cryo-EM samples were prepared by diluting a stock of TlpD SS1 (23 mg/ml) to a concentration of 0.46 mg/ml in a buffer consisting of 150 mM NaCl, 100 mM Tris pH 7 and 1 mM Tris(2-carboxyethyl)phosphine (TCEP). Samples were adsorbed onto freshly glow-discharged 300 mesh gold Quantifoil R1.2/1.3 grids (SPI Supplies, 4330G-CF) with PELCO easiGlow glow discharger and flash frozen in liquid ethane using either Vitrobot Mark IV dual-blotting plunger or Leica GP Climate controlled sample plunger with a controlled temperature and humidity. The grids were extensively screened with a 200kV Thermo Fisher Scientific Glacios electron microscope equipped with Falcon4i direct electron detector using the software SerialEM 4.0 for automated data collection. Typical acquisition parameters were as follows: dose rate of 4.88 e^-^/px/s in a counting mode, magnification of 130,000 times corresponding to the pixel size of 1.037 Å/px with a defocus range -0.5 to -2.5 μm (Extended Data Fig. 7-8). We collected 32 frames with 250 ms per frame exposure, a total exposure of 8 s and an accumulated dose of 39.04 e^-^. Cryo-EM grids displaying the best ice quality and particle distribution were used for data acquisition using a 300 kV Thermo Fisher Scientific Titan Krios G4 equipped with BioQuantum energy filter, and Gatan K3 and Falcon3 direct electron detectors. We acquired a total of 9,734 movies using software SerialEM 4.0 and the following acquisition parameters: magnification of 105,000 times corresponding to the pixel size of 0.820 Å/px with a defocus range from -0.8 μm to -2 μm. We collected 50 frames with an accumulated dose of 56.108 e^-^/A^2^.

### Cryo-EM structure determination

The structure of TlpD was calculated using cryoSPARC 4 ^73,74^. The following description outlines details of the protocol that yielded the final map deposited to the Electron Microscopy Data Bank (Fig 1f, Extended Data Fig. 7-8, Supplemental Table 2). 9,734 movies were patch motion- and patch CTF-corrected at the pixel size of 1.644 Å/px. Following inspection of pre-processed averaged micrographs, 206 out of 9,734 micrographs were discarded from further processing. We manually picked 2,000 particles from the selected micrographs to create 2D templates for automated particle picking. Particles were then automatically picked from a set of 9,528 curated and denoised micrographs using the template picker with templates representing a combination of different views of the complex. The template picker automatically selected 2,274,187 particle images. Selected particles were inspected and extracted from averaged micrographs with a box size of 180 pixels and the pixel size of 1.644 Å/px. Particle stacks were subsequently subjected to five rounds of 2D classification and averaging.

During the 2D analysis, artifacts and particles not converging to stable classes were removed from the data set. The resultant set of 528,846 particle images was then used for *ab-initio* reconstruction with four classes, followed by one round of heterogenous refinement. The three out of four classes representing different oligomeric forms of TlpD, namely monomers (31.6 % of particles), dimers (25.2%), and trimers (25.8%) of TlpD homodimers, were then further individually refined by homogenous and non-uniform refinement. Next, we minimized effects of continuous structural heterogeneity evidenced by the results of the 3D variability analysis by 3D flexible refinement (3D Flex) i.e. the deep-learning algorithm that employs graph neural networks for modeling flexible motions in biomolecules ^74^. In a following step, a set of 167,000 particles used to calculate a 3D map of the TlpD homodimer was subjected to 3D classification with six classes while employing C1 or C2 symmetry.

The 3D maps and correspondent particle stacks representing the two most populated classes from each run were combined (108,710,890 particle images corresponding to 65.1 % of all particles for a reconstruction obtained using the C1 symmetry, and 125,470 particle images corresponding to 75.2 % of all particles for a reconstruction representing the C2 symmetry) and subjected to an additional step of flexible refinement and reconstruction. To further improve map quality, we systematically tested several different sharpening methods, including tools relying on global Guinier plot-based sharpening, either automatic (PhenixAutoSharpen) or manual (CryoSPARC sharpening tool), local automatic Wiener filter-based methods (LocalDeblur and CryoSPARC Local Resolution Estimation/Filtering), and the convolutional neural network-based algorithm (DeepEMhancer). We also performed map phasing with Phenix Density Modification ^75^. The nominal resolution of TlpD structures calculated using the “gold standard” Fourier Shell Correlation (FSC) method according to the 0.143 criterion in cryoSPARC 4 ^73^ yielded 4.18 Å and 4.2 Å (Supplemental Data Table 2) maps for the C1 and C2 symmetries, respectively (Extended Data Fig. 7-8).

Interestingly, the FSC curves displayed a dip at approximately 9 Å resolution (Extended Data Fig. 8) that is not due to masking, as we thoroughly investigated this possibility by applying soft static and dynamic masking parameters. The FSC curves corresponding to the no mask and spherical mask reconstructions are nearly identical, indicating a well-behaved reconstruction with low background noise relative to the signal. The FSC curves obtained with tight and corrected masks are also nearly identical, indicating that the mask is not too tight and there is no overfitting. One possibility is that a dip at ∼9 Å resolution may reflect increased disorder at this and near spatial frequencies, as is often observed, for example, in membrane proteins stabilized by detergent micelles surrounding the protein. Interestingly, at low thresholds, the TlpD maps calculated with either C1 or C2 symmetry both revealed a belt-like structure wrapping around the MA at the base of CZB domains (Extended Data Fig. 8b). The feature likely represents associated lipid particles co-purified with TlpD, while providing an explanation for an observed dip in FSC curves (Extended Data Fig. 8a).

Another sharp dip in the corrected FSC curve indicates the resolution at which noise-substitution was automatically applied in cryoSPARC 4 ^73^. This procedure was designed to minimize effects of overfitting and resolution overestimation. The dip is an artifact of the boundary created at the spatial frequency where the phases were randomized versus where they were experimentally determined.

Additionally, the conical FSC (cFSC) plot, which reflects directional resolution anisotropy, yielded a cFAR value of 0.58, indicating sufficient Fourier sampling (Extended Data Fig. 8c). Consistent with this result, the angular distribution heat map confirmed nearly complete coverage of the angular space in the cryo-EM data (Extended Data Fig. 8e). To quantitatively assess the extent of sampling in Fourier space, we also monitored the Sampling Compensation Factor (SCF) ^68^ during structure calculations. The SCF value obtained for the final reconstruction is 0.922 (Extended Data Fig. 8f). The SCF values of 0.81 or higher are indicative of sufficient Fourier sampling ^68^. In addition, we monitored the relative signal (Extended Data Fig. 8g) and the high average relative signal amount within azimuth-elevation viewing regions (Extended Data Fig. 8g), as these two parameters are also sensitive to the extent of sampling in Fourier space ^69^. The algorithms compute cross-correlation coefficients between the two half maps as a function of viewing directions, where values of 0.5 and higher indicate that the contributing stack of particle images represents 2D projections of the complex that are accurately and uniformly distributed in orientation space. The cross-correlation coefficient values computed for the final reconstruction range from 0.8 to 1 across the entire orientation space, and from 0.81 to 0.84 across the highlighted viewing directions, both indicating sufficient Fourier sampling.

The intrinsic dynamics (Supplemental Movie 1) and heterogeneity (Extended Data Fig. 7) of TlpD samples precluded *de novo* model building directly from the map. We evidenced the continuous structural heterogeneity ^35^ of TlpD by employing the 3D Flex and 3D variability analyses, the fact further supported by the gradient in local resolution across the map (Extended Data Fig. 8d, Supplemental Movie 1). The initial model of TlpD was obtained by docking the *P*2_1_2_1_2_1_ X-ray structure of TlpD into the Coulomb density map using UCSF ChimeraX. The model was inspected and then iteratively refined by employing a combination of COOT 1.1 and Phenix 1.21 with imposed NCS, secondary structure and Ramachandran restraints. The atomic model obtained by refining the X-ray structure against the cryo-EM map overlays with the *P*2_1_2_1_2_1_ crystal structure, with a root mean square deviation (RMSD) of 1.13 Å calculated for 245 pruned Cα atom pairs (Extended Data Fig. 9g).

### TlpD Conservation and Phylogenetics

Filtration of the protein database generated by Perkins *et al.* in 2021 was performed to identify non-redundant CZB-containing proteins across diverse genera and within the order *Campylobacterales* ^35^. The sequence of *H. pylori* TlpD J99 was also used as a search query in the software Geneious Prime 2025 with default cutoffs, searching through all notated *H. pylori* sequences as of August 2025. Sequences were aligned using MUSCLE and phylogenetic trees were constructed with phyloT (Fig. 2e-f, Supplemental Data 1).

## Acknowledgments

Funding for this work was provided by NIAID under award numbers 1K99AI148587 and 4R00AI148587-03, the Stanley Adler Fund, the EschLEAD program, and startup funding from Washington State University to AB, and a start-up grant from Rutgers University to A.W.K. This research used resources of the Advanced Light Source, through beamline 5.0.2., a U.S. DOE Office of Science User Facility under contract no. DE-AC02-05CH11231. We thank Melissa Chambers for her assistance with EM data collection, which was conducted in part at the Center for Structural Biology Cryo-EM Facility at Vanderbilt University.

## Author Contributions

K.F. performed crystallographic data processing, refinements, and analyses. M.D. conducted negative-stain EM experiments. A.S. and R.K. assisted with solution methods for the P1 TlpD structure. K.K., E.S., and M.C. carried out cloning, purification, and crystallization of TlpD. S.J.R. supervised the design of TlpD molecular and structural studies and contributed to data analysis and interpretation. A.W.K. performed cryo-EM experiments, structure determination, model building, and analysis. A.B. carried out TlpD crystal harvesting and heavy atom soaks, crystallographic data collection, structure determination, model building, and analysis. All authors contributed to writing and revising the manuscript.

## Data availability

All data and materials that are necessary to evaluate the conclusions of the paper are available from the corresponding author(s) upon reasonable request. Crystallographic data have been deposited in the Protein Data Bank (PDB) under the identifiers 10ea for the *P*1 TlpD structure (zinc-disrupted and zinc-strained homodimers) and 10DZ for the *P*2_1_2_1_2_1_ TlpD structure (zinc-intact homodimer). The cryo-EM map of TlpD has been deposited in the Electron Microscopy Data Bank (EMDB) under the accession code EMD-78945 (zinc-intact homodimer).

## Extended Data Figures

**Extended Data Fig. 1.**
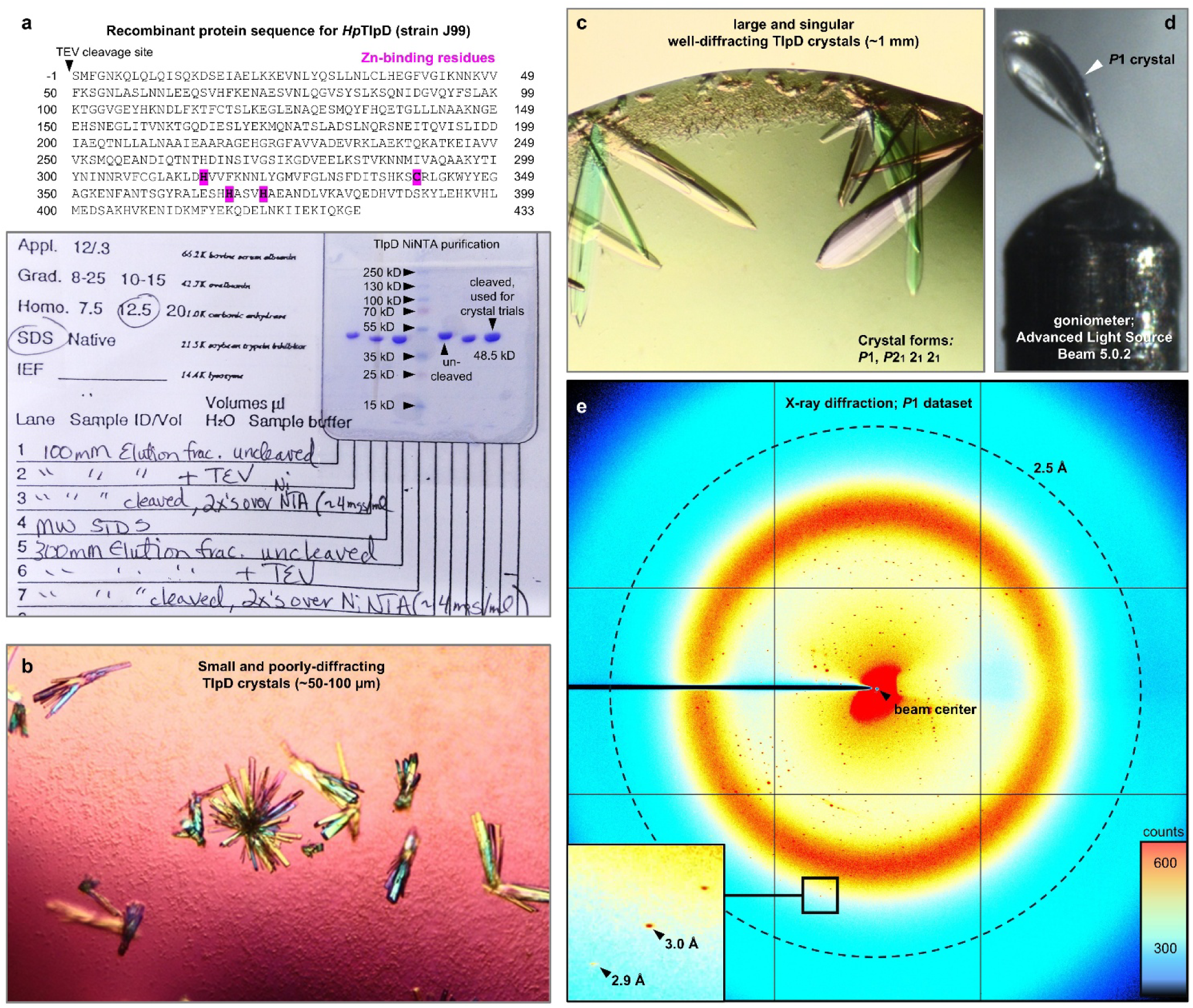
Crystallization of *Helicobacter pylori* TlpD strain J99. **a.** Full amino acid sequence of the crystallized protein. *H. pylori* TlpD from strain J99 was expressed with an N-terminal 6x His-tag containing a TEV cleavage site; an N-terminal Ser remains following TEV cleavage. Zinc-binding residues are highlighted in pink. Shown bottom are steps from Ni-NTA purification and TEV-cleavage for the protein used for crystallography. **b.** Shown is a representative image of the typical small clusters of poorly-diffracting crystals that TlpD readily forms in a variety of PEG and ammonium sulfate-based crystallization conditions. **c.** An image of the single drop in which large and well-diffracting TlpD crystals grew after approximately 2 years that yielded the datasets in this study. **d.** An image of the *P*1 crystal (harvested from **c**) that yielded the initial solution. The crystal was flash frozen and x-ray diffraction data collected under cryo conditions. **e.** A representative diffraction image from the crystal in **d**.

**Extended Data Fig. 2.**
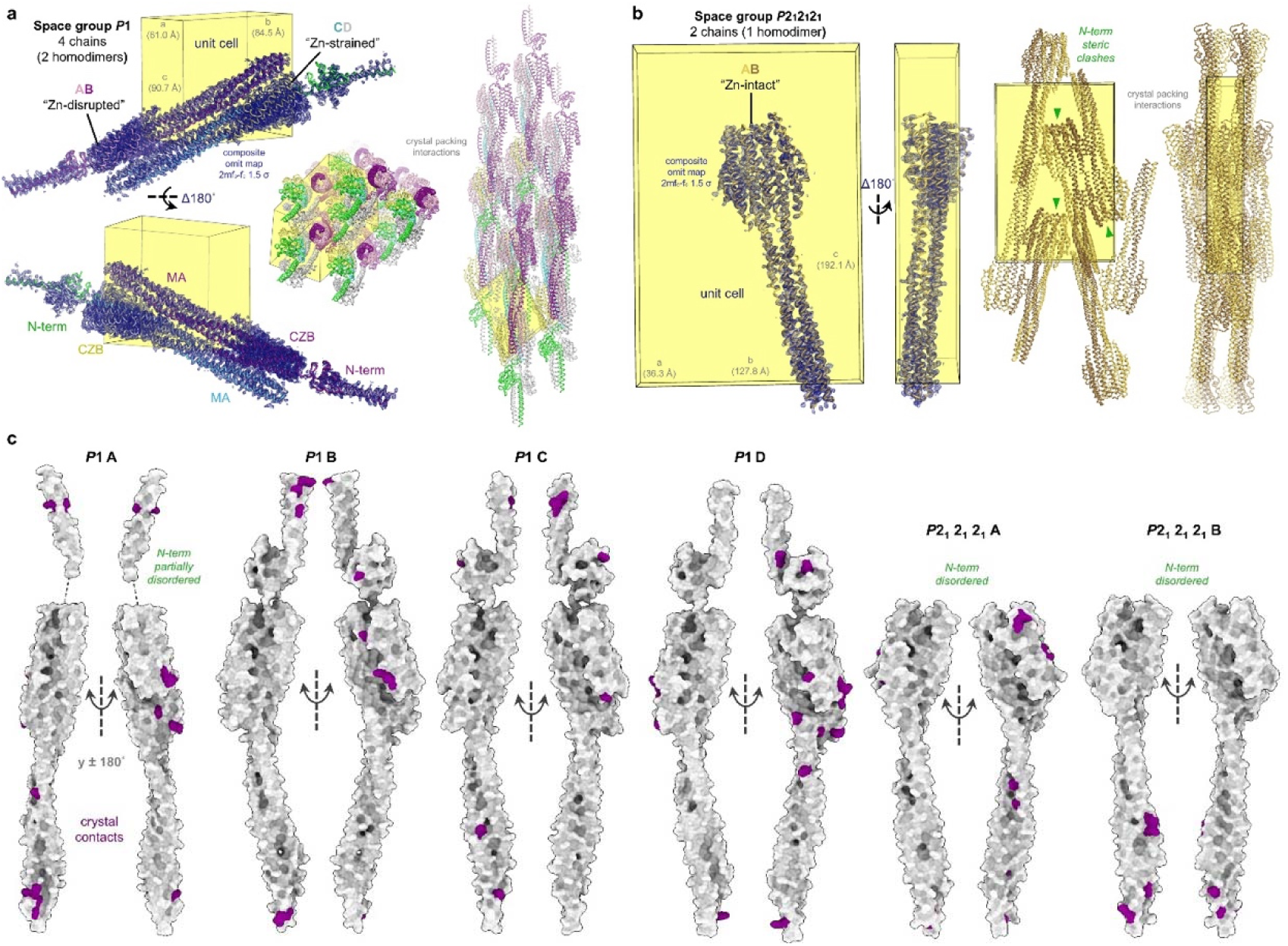
Solution of *Helicobacter pylori* TlpD J99 in two crystal forms. **a.** Shown are the crystal arrangements for the *P*1 crystal structure of TlpD, consisting of two homodimers within the asymmetric unit that we refer to as the conformations “zinc-disrupted” (chains AB, light and dark pink, respectively) and “zinc-strained” (chains CD, colored by domain, as indicated, and gray, respectively). The N-terminal PAS is green, the MA domain is light blue, and the CZB domain is yellow. Electron density (dark blue mesh, 2fo-fc) is from a composite omit map generated with 5% of the atoms removed. For all chains, the N-terminal PAS domain is poorly ordered and exhibits weak electron density. **b.** The crystal arrangements for the *P*2_1_2_1_2_1_ TlpD crystal structure composed of Chains AB (gold and brown), which we refer to as “zinc-intact.” Arrows (green) note crystal contact interactions that pose steric clashes for a folded N-terminal region, and hence this region of the structure is disordered and unresolved in this crystal form. **c.** Residues involved in crystal contacts for each chain are colored in magenta.

**Extended Data Fig. 3.**
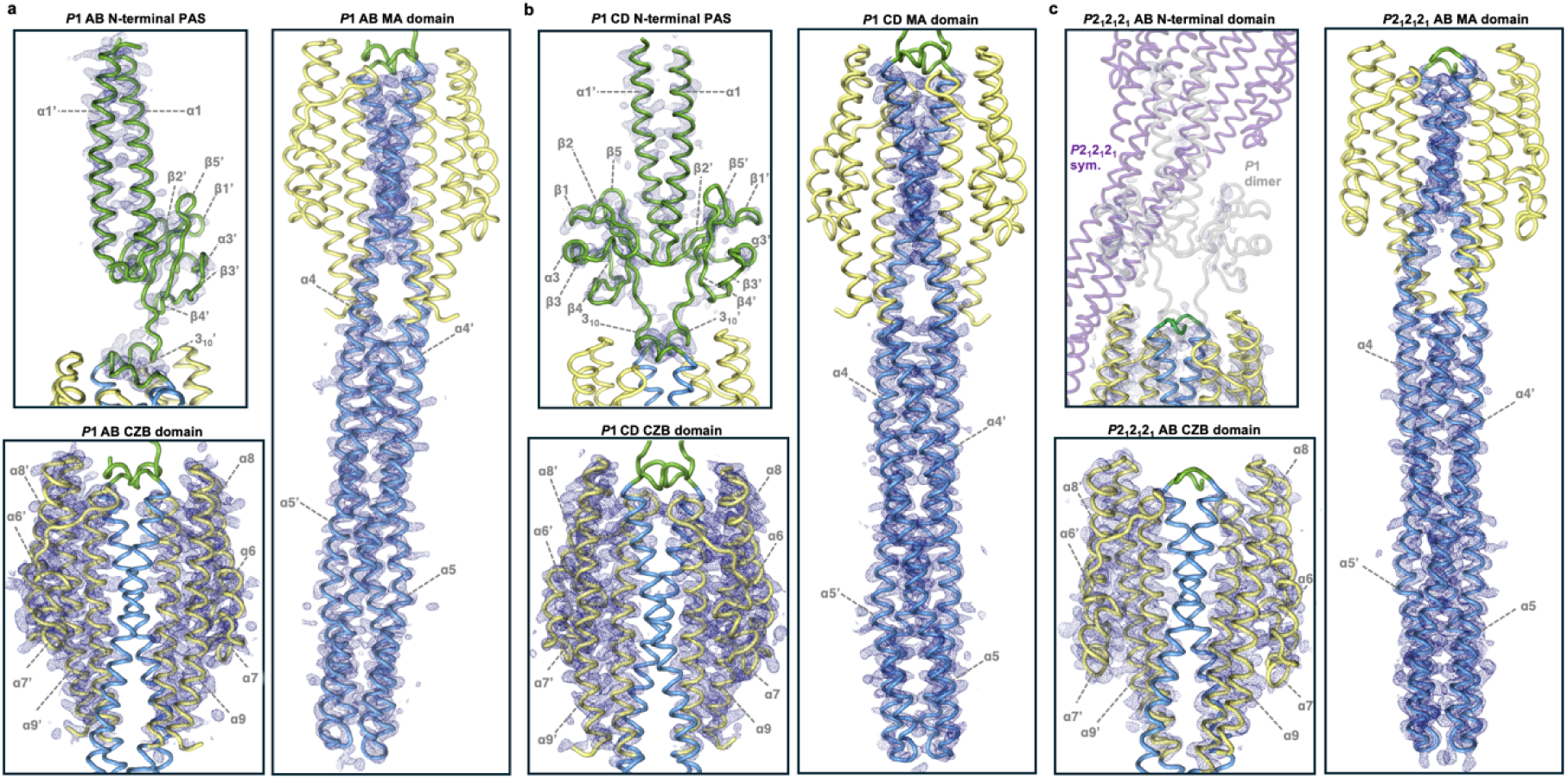
Electron density quality of the TlpD homodimers. **a-c**. Electron density for each TlpD homodimer is shown for different regions of the protein, as indicated. Dark blue mesh represents 2fo-fc omit map density at 1 σ calculated with 5% of atoms missing. with the protein colored by domain: green for N-terminal PAS, light blue for the MA domain, and yellow for the CZB domain. For the *P*2_1_ 2_1_ 2_1_ N-terminal PAS, a symmetry mate is shown in purple, and the N-terminal region of the *P*1 CD homodimer is modeled in gray, to illustrate the steric clash that prevents a folded N-terminus in this structure.

**Extended Data Fig. 4.**
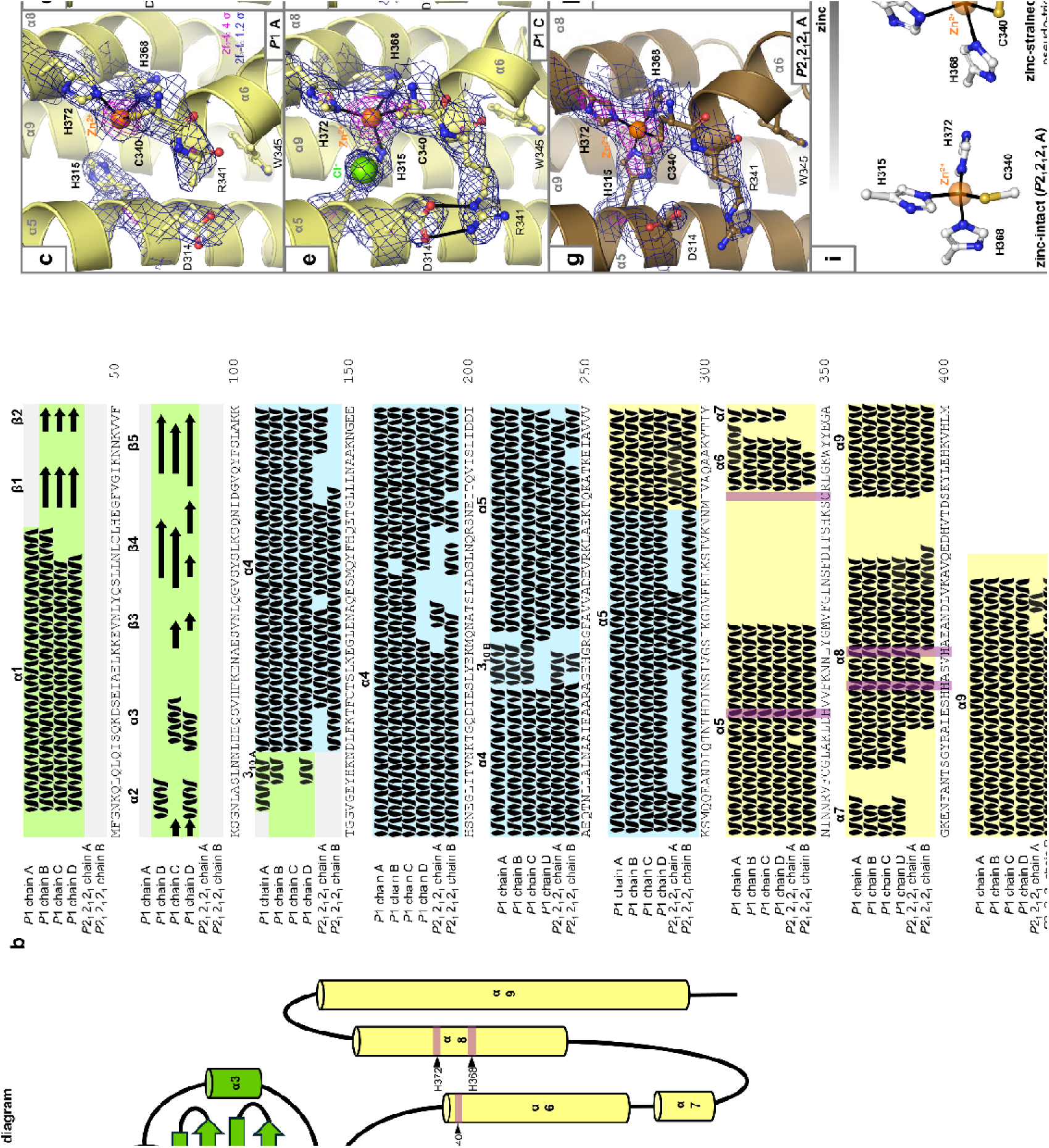
Topology and zinc-binding state of TlpD crystal structures. **a.** Consensus topology diagram based on the secondary structure elements observed in the crystal structures, with helices denoted as cylinders and strands as arrows and loops as lines; lengths of helices and strands are proportional to their size in the structures. Domain regions are colored as in Fig. 1, with zinc-binding residues noted in magenta. **b.** Secondary structure assignments for each TlpD crystal structure chain based on DSSP. **c-h.** Electron density for the zinc-binding sites of the TlpD crystal structures, as indicated. Halogens are noted. **i.** Summary of key zinc-binding site features for the zinc-intact, zinc-strained, and zinc-disrupted conformations.

**Extended Data Fig. 5.**
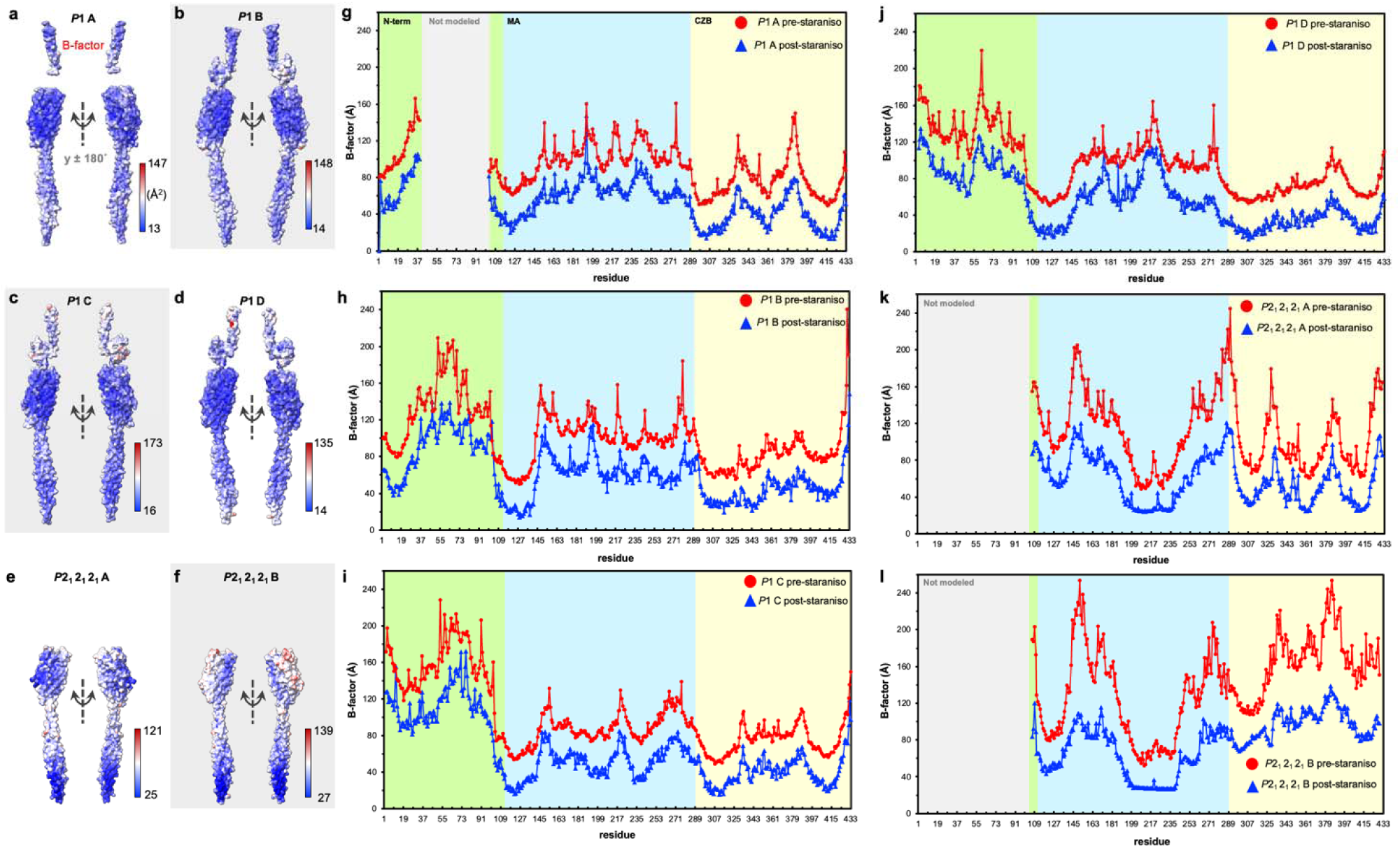
Analysis of crystallographic B-factors. **a-f**. Each TlpD chain is shown with molecular surface representation colored by B-factor, as indicated. **g-l**. Plots showing B-factor per residue before (red circles) and after (blue triangles) Staraniso data processing. Domain designations are colored as in Fig. 1a, or gray for regions not modeled.

**Extended Data Fig. 6.**
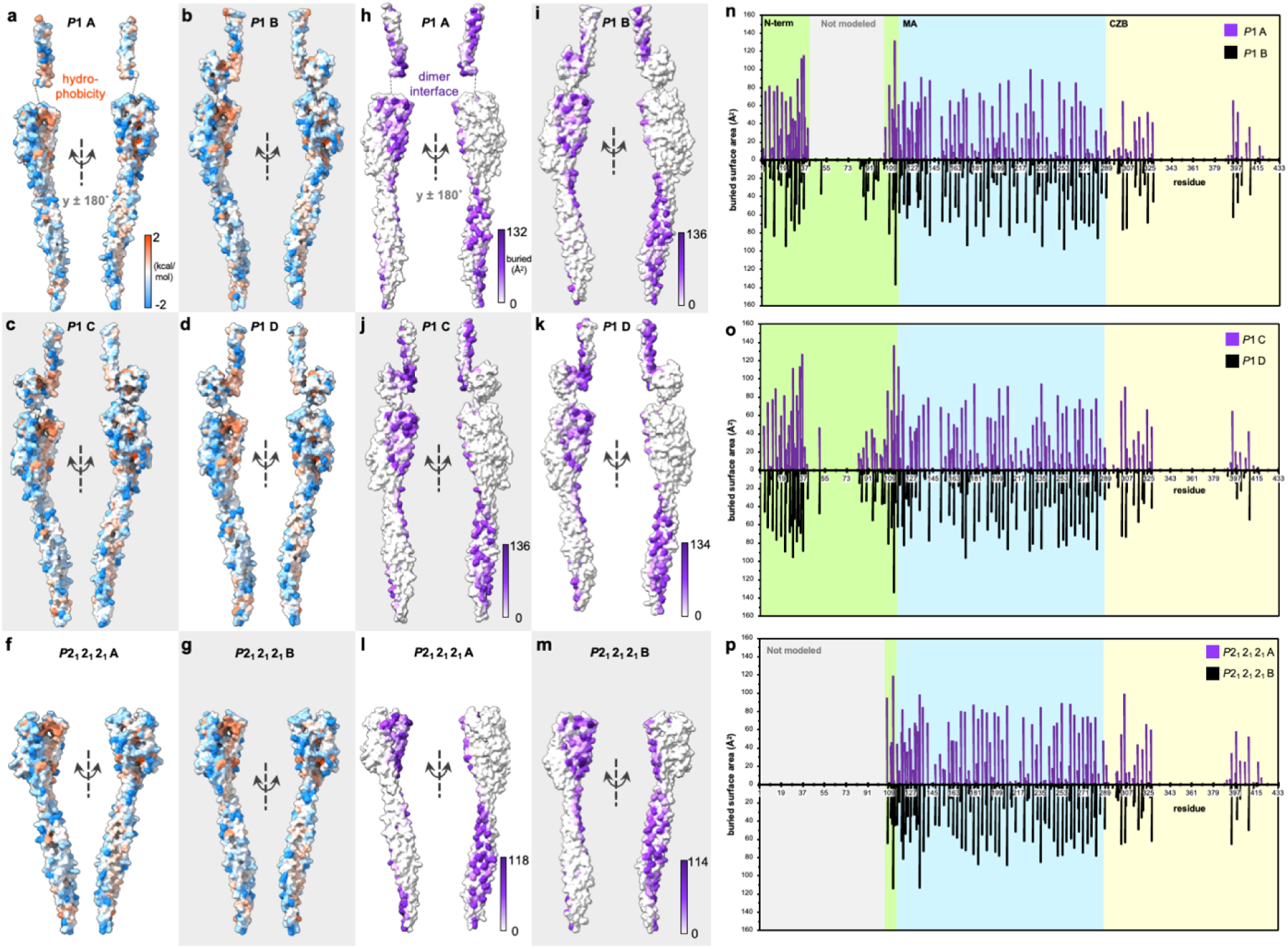
The homodimer interface of TlpD. **a-d.** *P*1 chains colored by hydrophobicity using the Wimley-White hydrophobicity scale, which is based on the free energy associated with transitioning a peptide from an aqueous environment to a hydrophobic environment in units of kcal/mol. **e-f.** *P*2_1_2_1_2_1_ chains colored by hydrophobicity using the Wimley-White hydrophobicity scale. **g-l.** TlpD homodimer interface of all chains from the *P*1 and *P*2_1_2_1_2_1_ crystal structures, as indicated. **m-o.** Buried surface area per residue, for the *P*1 TlpD AB homodimer, CD homodimer, and *P*2_1_2_1_2_1_ AB homodimer, as indicated. Plot regions are colored by domain as in Fig. 1a, or gray for regions not modeled.

**Extended Data Fig. 7.**
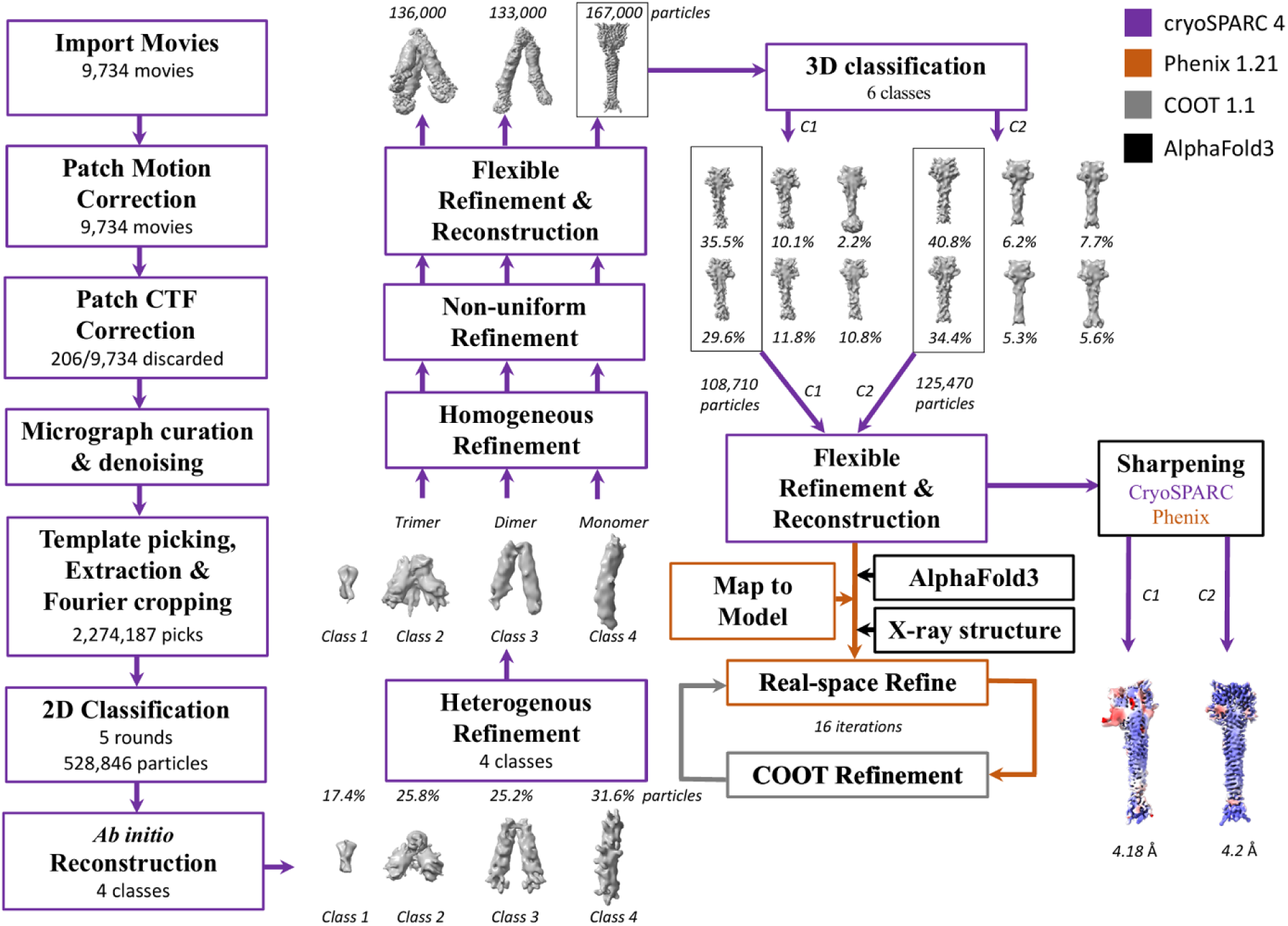
The SPA workflow employed for cryo-EM structure calculations. We used a combination of cryoSPARC 4, Phenix 1.21, COOT 1.1 and Alphafold 3 for image processing and structure calculations of TlpD. Steps completed in each program are color-coded according to the figure legend. For details, please see the Methods section.

**Extended Data Fig. 8.**
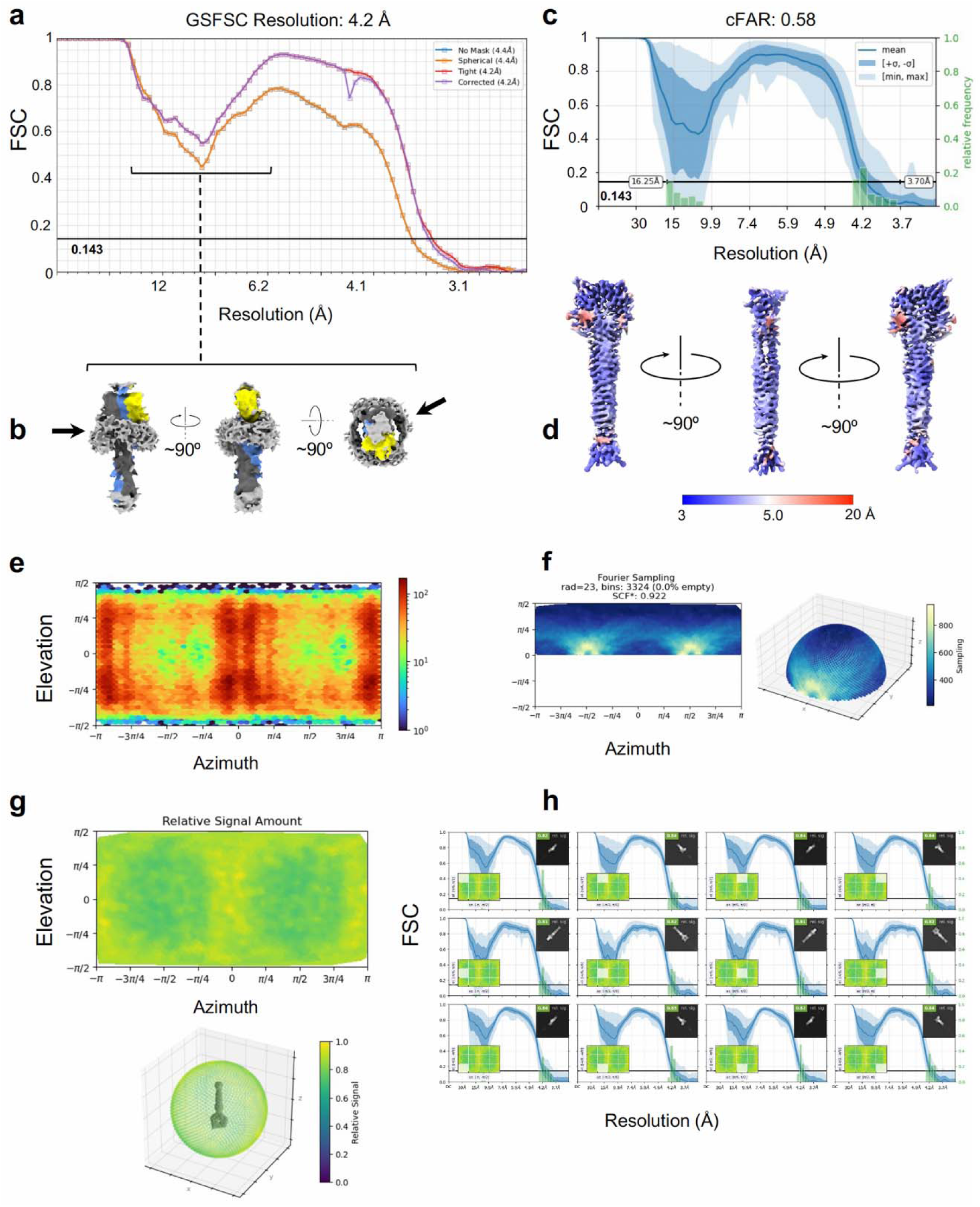
Evaluation of the TlpD Coulomb map resolution and the extent of Fourier space sampling. **a.** The nominal resolution of TlpD calculated using the “gold standard” Fourier Shell Correlation (FSC) method according to the 0.143 criterion in cryoSPARC 4 is 4.2 Å. The FSC curves display a dip at approximately 9 Å resolution, which is not due to masking. **b.** At lower thresholds, the TlpD map revealed a belt-like structure wrapping around the MA at the base of CZB domains possibly indicating the presence of associated lipid particles co-purified with TlpD, while providing an explanation for observed dip in FSC curves. **c.** The conical FSC (cFSC) plot reflecting directional resolution anisotropy. The cFAR value of 0.58 indicates sufficient Fourier sampling. **d.** Local resolution values mapped onto the Coulomb map of TlpD. The gradient of local resolutions likely reflects the intrinsic dynamics of the complex. **e.** The angular distribution heat map confirms the complete coverage of the angular space in the cryo-EM data. **f**. Sampling Compensation Factor (SCF) during structure calculations. The SCF value obtained for the final reconstruction is 0.922. The SCF values of 0.81 or higher are indicative of sufficient Fourier sampling. **g.** The relative signal and **h.** the high average relative signal amount within azimuth-elevation viewing regions. The cross-correlation coefficient values computed for the final reconstruction range from 0.8 to 1 across the entire orientation space, and from 0.81 to 0.84 across the highlighted viewing directions, both indicating sufficient Fourier sampling.

**Extended Data Fig. 9.**
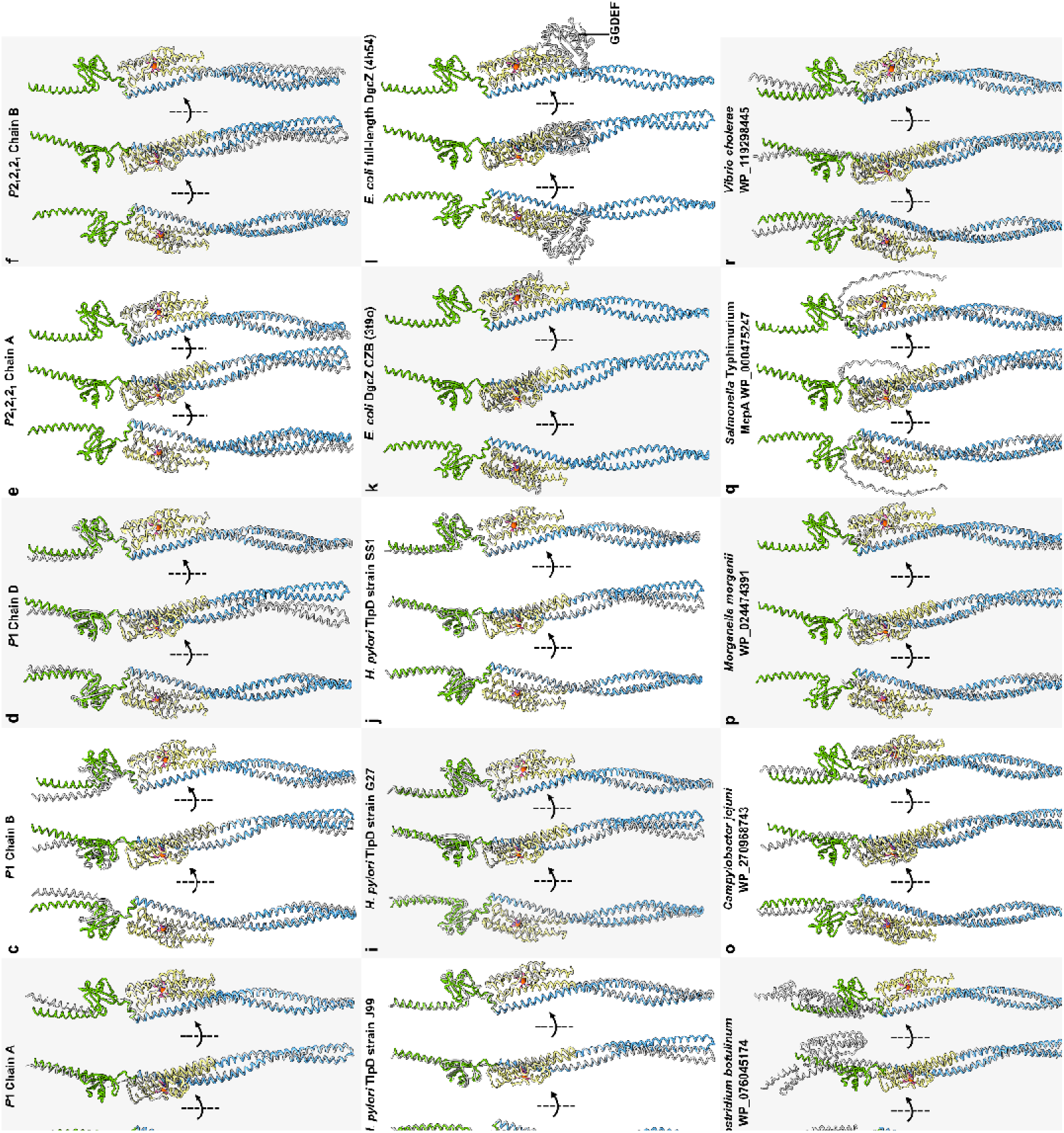
Structural overlays with TlpD. **a**. Reference chain C from the TlpD *P*1 crystal structure in three orientations, colored as in Fig. 1a, with the zinc ion shown in orange as an enlarged sphere for clarity**. b-d.** Chains A, B and D (white) from the TlpD *P*1 crystal structure are shown overlaid onto reference chain C. **e-f.** Chains A and B from the *P*2_1_2_1_2_1_ crystal structure are shown (white) overlaid onto reference chain C. **g.** Overlay with the cryo-EM TlpD structure from strain SS1. **h-j.** chain A (white) from Alphafold 3 models of TlpD from *H. pylori* strain J99, G27 and SS1 respectively, were overlaid onto reference chain C. AlphaFold 3 models were generated as homodimers with chain A used for the overlays. **k-l.** Chain A from the crystal structures of the *E. coli* DgcZ CZB fragment, and full-length structure of a C52A mutant, PDB ascension codes 3t9o and 4h54, respectively, are shown overlaid onto reference chain C. **m-r.** Alphafold 3 models of six different TlpD homologs (white) from various bacterial families are shown overlaid onto reference chain C. AlphaFold 3 models were generated as homodimers with chain A used for the overlays. Residues 333-352 of the CZB domain in chain C were used for the alignments. **s.** Overlay of TlpD hairpin tips, using residues indicated. See also Supplemental Table 3 for Cα RMSD values.

**Extended Data Fig. 10.**
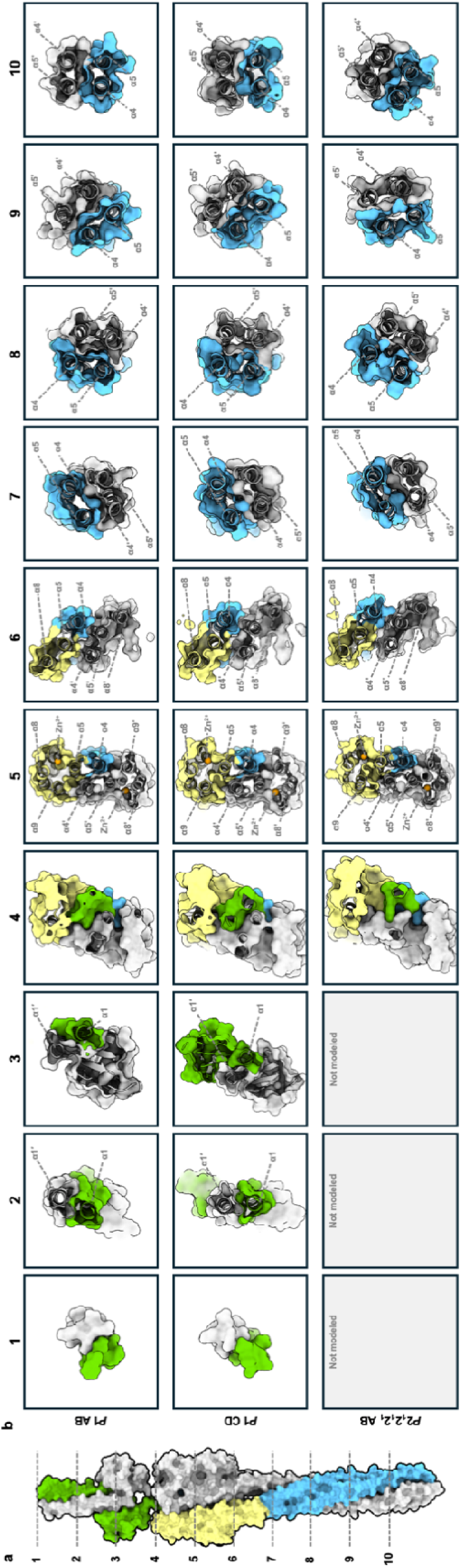
TlpD structures comparedthrough the coiled-coil axis. **a.** Positions of cross- sectional slices taken at 24 Å intervals for each TlpD structure along the central coiled-coil axis are indicated with dashed lines. **b.** For each indicated position (1-10), top-down views are shown of the corresponding cross section in panel a. Each panel represents a 24 Å-thick slab, revealing changes in oligomeric architecture and interfaces along the length of the protein. Chains A and C of the *P*1 structure, and A of the *P*2_1_2_1_2_1_ structure are in Fig. 1a, with partnercolored by domain as chains in white.Comparisons across panels 8-10 clearly show the increased supercoiling at the hairpin tip for the *P*1 chains relative to the *P*2_1_2_1_2_1_ chains.

## Supplemental Tables

**Supplemental Table 1:** Summary of crystallographic statistics.

| Protein, PDB code | *H. pylori* TlpD<sub>J99</sub>, PDB: 10ea | *H. pylori* TlpD<sub>J99</sub>, PDB: 10dz |
| --- | --- | --- |
| Space Group | *P* 1 | *P* 2<sub>1</sub> 2<sub>1</sub> 2<sub>1</sub> |
| Cell dimensions and angle<br>(a, b, c, $\alpha$, $\beta$, $\gamma$) (Å, °) | 61.0, 84.5, 90.7, 92.2, 90.4,<br>103.7 | 36.3, 127.8, 192.1, 90, 90, 90 |
| Resolution (Å)<sup>a</sup> | 49.2-2.60 (2.67-2.60) | 33.94-2.92 (3.07-2.92) |
| Resolution limit (each axis, Å) | 2.45, 2.90, 3.28 | 3.07, 3.07, 2.92 |
| Completeness (%)<sup>a</sup> | 67.33 (10.47) | 97.9 (92.7) |
| Total reflections | 508149 (968) | 155943 (8397) |
| Unique reflections | 131548 (419) | 27093 (1916) |
| Average $I/\sigma$<sup>a</sup> | 2.29 (9.73) | 4.18 (2.34) |
| R<sub>merge</sub><sup>a</sup> | 0.66 (0.62) | 0.1801 (5.967) |
| CC<sub>1/2</sub><sup>a</sup> | 0.85 (0.15) | 0.997 (0.018) |
| R<sub>work</sub> (%)<sup>b</sup> | 22.6 | 20.3 |
| R<sub>free</sub> (%)<sup>b</sup> | 29.2 | 29.5 |
| Ramachandran favored,<br>allowed, outliers (%) | 95.3, 4.1, 0.6 | 84.8, 12.6, 2.64 |
| Non-hydrogen atoms | 12589 | 4979 |
| Solvent atoms | 37 | 9 |
| Protein chains, residues | 4, 1662 | 2, 648 |
| Average B-factor of protein<br>atoms (Å<sup>2</sup>) | 59.7 | 89.5 |
| Average B-factor of solvent atoms ($\text{\AA}^2$) | 27 | 32 |
| rms bond lengths ($\text{\AA}$) | 0.017 | 0.008 |
| rms bond angles ($^\circ$) | 1.57 | 1.04 |
| TLS groups | 13 | 9 |
| Molprobity clash score, percentile | 21.3, 75<sup>th</sup> | 11.7, 97<sup>th</sup> |

<sup>a</sup> Values in parentheses indicate statistics for the highest resolution shell.
<sup>b</sup> Only resolution out to 3.30 $\text{\AA}$ was used for refinement.

**Supplemental Table 2.** Cryo-EM data collection.

| TlpD<br>(EMD-78945 [https://www.ebi.ac.uk/emdb/EMD-78945]) |  |
| --- | --- |
| **Data collection and processing** |  |
| Magnification | 105,000x |
| Voltage (kV) | 300 |
| Electron exposure ($\text{e}/\text{\AA}^2$) | 56.1 |
| Defocus range ($\mu\text{m}$) | 0.8-2.0 |
| Pixel size ($\text{\AA}$) | 1.64 |
| Symmetry imposed | C2 |
| Initial particle images (no.) | 2,274,187 |
| Final particle images (no.) | 125,000 |
| Map resolution ($\text{\AA}$) | 4.2 |
| FSC threshold | 0.143 |
| Map resolution range ($\text{\AA}$) | 3-15 |

**Supplemental Table 3.**
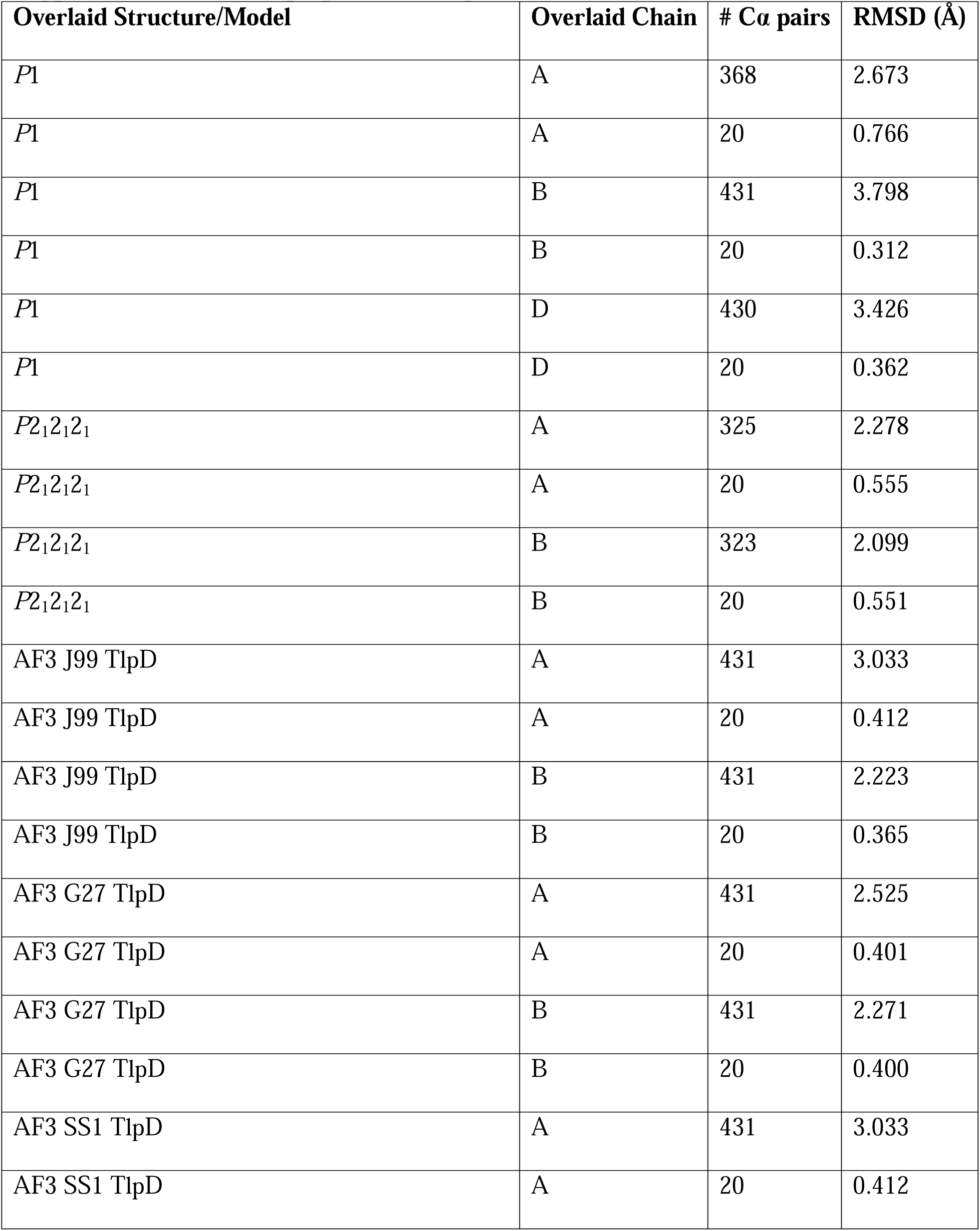
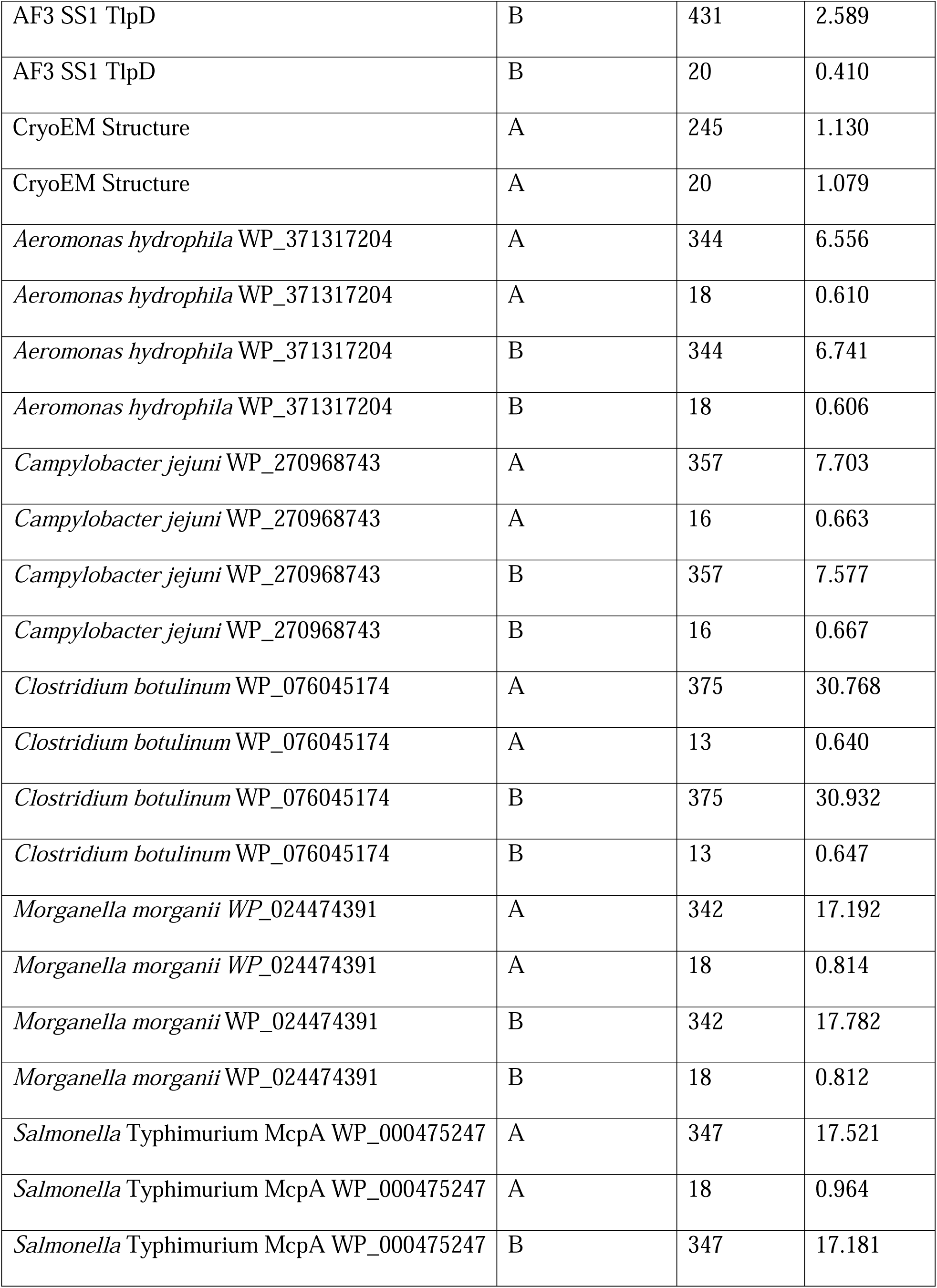
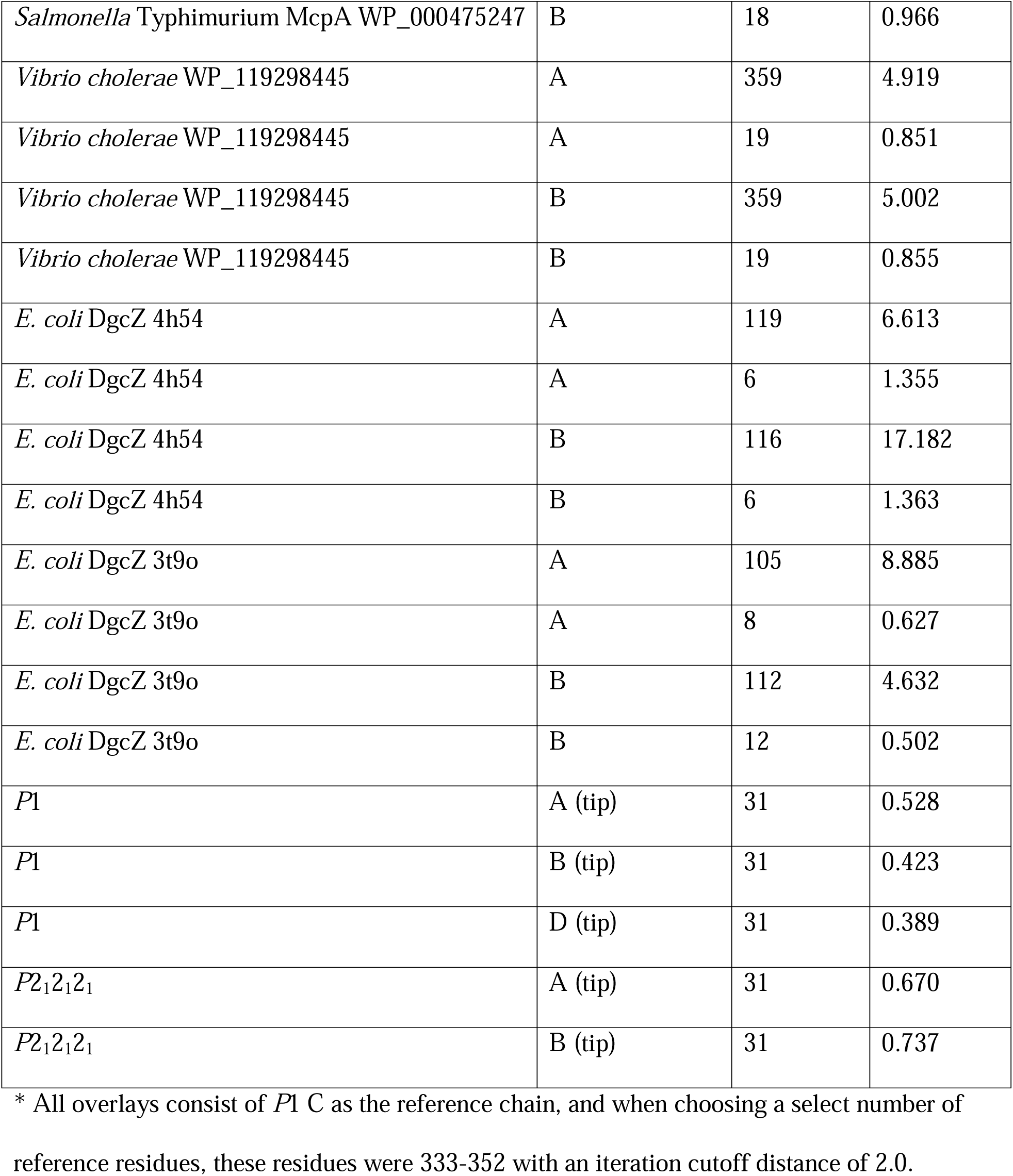
Superposition of TlpD structures and models.

## Supplemental Movies

**Supplemental Movie 1.** Structural features of TlpD. Shown is a video of the major structural features of the TlpD homodimer, focusing on the zinc-strained conformation (Chains CD of the *P*1 crystal structure). The domains, key structural elements and residues, and electron density is shown, as well as a morph between the zinc-strained and zinc-disrupted conformations. We employed 3D flexible refinement (3D Flex), which uses graph neural networks to model flexible motions in biomolecules. Analysis of the cryo-EM data using 3D Flex revealed coupled motions of the CZBs and MA in solution, including coordinated rearrangements of the CZBs associated with gear-like rotation of the interlocked helices around the central coiled-coil axis. We used 3D variability analysis based on principal component analysis (PCA) to quantitatively assess structural heterogeneity in the cryo-EM data acquired for TlpD. The analysis revealed continuous structural heterogeneity rather than discrete conformational states, consistent with the 3D flexible refinement results. Variability within the CZB domains is coupled to rotation of individual α-helices within the MA domains and conformational changes at their tips. Viewable at: https://youtu.be/0jsKlSzdjgU.

## Supplemental Data

**Supplemental Data 1. TlpD orthologue sequences and Alphafold 3 model database.** Sequences, organism, and other metadata for TlpD orthologs are provided, as well as coordinates for homodimer models generated using Alphafold 3. Data accessible at the following public repository: https://public.vetmed.wsu.edu/Baylink/Franco-et-al-biorxiv-2026

## References

1. Lupas, A. N. & Bassler, J. Coiled Coils - A Model System for the 21st Century. Trends Biochem Sci 42, 130–140 (2017).

2. Woolfson, D. N. Understanding a protein fold: The physics, chemistry, and biology of α-helical coiled coils. The Journal of Biological Chemistry 299, 104579 (2023).

3. Truebestein, L. & Leonard, T. A. Coiled-coils: The long and short of it. Bioessays 38, 903– 916 (2016).

4. Hazelbauer, G. L., Falke, J. J. & Parkinson, J. S. Bacterial chemoreceptors: high-performance signaling in networked arrays. Trends in Biochemical Sciences 33, 9–19 (2008).

5. Perrot, R. & Eyer, J. Neuronal intermediate filaments and neurodegenerative disorders. Brain Res Bull 80, 282–295 (2009).

6. Munro, S. The Golgin Coiled-Coil Proteins of the Golgi Apparatus. Cold Spring Harb Perspect Biol 3, a005256 (2011).

7. Kon, T. et al. Helix sliding in the stalk coiled coil of dynein couples ATPase and microtubule binding. Nat Struct Mol Biol 16, 325–333 (2009).

8. Walshaw, J. & Woolfson, D. N. Extended knobs-into-holes packing in classical and complex coiled-coil assemblies. Journal of Structural Biology 144, 349–361 (2003).

9. Alexander, R. P. & Zhulin, I. B. Evolutionary genomics reveals conserved structural determinants of signaling and adaptation in microbial chemoreceptors. Proceedings of the National Academy of Sciences 104, 2885–2890 (2007).

10. Lazzari, E. & Meroni, G. TRIM32 ubiquitin E3 ligase, one enzyme for several pathologies: From muscular dystrophy to tumours. Int J Biochem Cell Biol 79, 469–477 (2016).

11. Davies, K. E. & Nowak, K. J. Molecular mechanisms of muscular dystrophies: old and new players. Nat Rev Mol Cell Biol 7, 762–773 (2006).

12. Kon, T. et al. The 2.8 Å crystal structure of the dynein motor domain. Nature 484, 345–350 (2012).

13. Schofield, A. V. & Bernard, O. Rho-associated coiled-coil kinase (ROCK) signaling and disease. Crit Rev Biochem Mol Biol 48, 301–316 (2013).

14. Parkinson, J. S., Hazelbauer, G. L. & Falke, J. J. Signaling and sensory adaptation in Escherichia coli chemoreceptors: 2015 update. Trends Microbiol 23, 257–266 (2015).

15. Yang, W. et al. In Situ Conformational Changes of the Escherichia coli Serine Chemoreceptor in Different Signaling States. mBio 10, e00973–19 (2019).

16. Flack, C. E. & Parkinson, J. S. Bacterial Chemoreceptors Transmit Stimulus Signals Through Coupled Entropic Switches. Journal of Molecular Biology 438, 169614 (2026).

17. Ortega, Á., Zhulin, I. B. & Krell, T. Sensory Repertoire of Bacterial Chemoreceptors. Microbiology and Molecular Biology Reviews 81, e00033–17 (2017).

18. Gumerov, V. M., Ulrich, L. E. & Zhulin, I. B. MiST 4.0: a new release of the microbial signal transduction database, now with a metagenomic component. Nucleic Acids Res 52, D647–D653 (2023).

19. Collins, K. D., Lacal, J. & Ottemann, K. M. Internal Sense of Direction: Sensing and Signaling from Cytoplasmic Chemoreceptors. Microbiology and Molecular Biology Reviews 78, 672–684 (2014).

20. Zhou, B., Szymanski, C. M. & Baylink, A. Bacterial chemotaxis in human diseases. Trends in Microbiology 10.1016/j.tim.2022.10.007 (2022) doi:10.1016/j.tim.2022.10.007.

21. Keegstra, J. M., Carrara, F. & Stocker, R. The ecological roles of bacterial chemotaxis. Nat Rev Microbiol 1–14 (2022) doi:10.1038/s41579-022-00709-w.

22. Matilla, M. A. & Krell, T. The effect of bacterial chemotaxis on host infection and pathogenicity. FEMS Microbiol Rev 42, (2018).

23. Schavemaker, P. E. & Lynch, M. Flagellar energy costs across the tree of life. eLife 11, e77266 (2022).

24. Matilla, M. A., Gavira, J. A. & Krell, T. Accessing nutrients as the primary benefit arising from chemotaxis. Current Opinion in Microbiology 75, 102358 (2023).

25. Matilla, M. A. & Krell, T. Chemoreceptor-based signal sensing. Current Opinion in Biotechnology 45, 8–14 (2017).

26. Flack, C. E. & Parkinson, J. S. Bacterial Chemoreceptors Transmit Stimulus Signals Through Coupled Entropic Switches. Journal of Molecular Biology 438, 169614 (2026).

27. Bi, S., Pollard, A. M., Yang, Y., Jin, F. & Sourjik, V. Engineering Hybrid Chemotaxis Receptors in Bacteria. ACS Synth. Biol. 5, 989–1001 (2016).

28. Gholami, A., Mohkam, M., Soleimanian, S., Sadraeian, M. & Lauto, A. Bacterial nanotechnology as a paradigm in targeted cancer therapeutic delivery and immunotherapy. Microsyst Nanoeng 10, 113 (2024).

29. Collins, K. D. et al. The Helicobacter pylori CZB Cytoplasmic Chemoreceptor TlpD Forms an Autonomous Polar Chemotaxis Signaling Complex That Mediates a Tactic Response to Oxidative Stress. Journal of Bacteriology 198, 1563–1575 (2016).

30. Behrens, W. et al. Localisation and protein-protein interactions of the Helicobacter pylori taxis sensor TlpD and their connection to metabolic functions. Sci Rep 6, 23582 (2016).

31. Rolig, A. S., Shanks, J., Carter, J. E. & Ottemann, K. M. Helicobacter pylori Requires TlpD-Driven Chemotaxis To Proliferate in the Antrum. Infect Immun 80, 3713–3720 (2012).

32. Huang, J. Y., Sweeney, E. G., Guillemin, K. & Amieva, M. R. Multiple Acid Sensors Control Helicobacter pylori Colonization of the Stomach. PLOS Pathogens 13, e1006118 (2017).

33. Cooper, K. G. et al. HilD-regulated chemotaxis proteins contribute to Salmonella Typhimurium colonization in the gut. mBio 16, e00390–25.

34. Perkins, A., Tudorica, D. A., Amieva, M. R., Remington, S. J. & Guillemin, K. Helicobacter pylori senses bleach (HOCl) as a chemoattractant using a cytosolic chemoreceptor. PLOS Biology 17, e3000395 (2019).

35. Perkins, A. et al. A Bacterial Inflammation Sensor Regulates c-di-GMP Signaling, Adhesion, and Biofilm Formation. mBio 12, (2021).

36. Zähringer, F., Lacanna, E., Jenal, U., Schirmer, T. & Boehm, A. Structure and Signaling Mechanism of a Zinc-Sensory Diguanylate Cyclase. Structure 21, 1149–1157 (2013).

37. Draper, J., Karplus, K. & Ottemann, K. M. Identification of a chemoreceptor zinc-binding domain common to cytoplasmic bacterial chemoreceptors. J Bacteriol 193, 4338–4345 (2011).

38. Zumwalt, L., Perkins, A. & Ogba, O. M. Mechanism and Chemoselectivity for HOCl-Mediated Oxidation of Zinc-Bound Thiolates. ChemPhysChem 21, 2384–2387 (2020).

39. Franco, K., Cooper, K. G., Shin, C. H., Steele-Mortimer, O. & Baylink, A. Zn2+ as a secondary messenger for exogenous redox potential sensed through Chemosensory Zinc-Binding (CZB) protein domains. 2026.07.03.735341 Preprint at 10.64898/2026.07.03.735341 (2026).

40. Glenn, S. J. et al. Bacterial vampirism mediated through taxis to serum. eLife 12, RP93178 (2024).

41. Boyd, A., Kendall, K. & Simon, M. I. Structure of the serine chemoreceptor in Escherichia coli. Nature 301, 623–626 (1983).

42. Pollard, A. M., Bilwes, A. M. & Crane, B. R. The structure of a soluble chemoreceptor suggests a mechanism for propagating conformational signals. Biochemistry 48, 1936–1944 (2009).

43. Cassidy, C. K. et al. CryoEM and computer simulations reveal a novel kinase conformational switch in bacterial chemotaxis signaling. eLife 4, e08419 (2015).

44. Ashkenazy, H. et al. ConSurf 2016: an improved methodology to estimate and visualize evolutionary conservation in macromolecules. Nucleic Acids Res 44, W344–350 (2016).

45. Burkhard, P., Stetefeld, J. & Strelkov, S. V. Coiled coils: a highly versatile protein folding motif. Trends Cell Biol 11, 82–88 (2001).

46. Lopez-Magaña, R. & Ottemann, K. M. The Helicobacter pylori TlpD cytoplasmic chemoreceptor requires an intact C-terminus for polar localization and function. J Bacteriol 208, e0039425 (2026).

47. Briegel, A. et al. Structure of bacterial cytoplasmic chemoreceptor arrays and implications for chemotactic signaling. eLife 3, e02151 (2014).

48. Moi, D. et al. Structural phylogenetics unravels the evolutionary diversification of communication systems in gram-positive bacteria and their viruses. Nat Struct Mol Biol 32, 2492–2502 (2025).

49. Abramson, J. et al. Accurate structure prediction of biomolecular interactions with AlphaFold 3. Nature 630, 493–500 (2024).

50. Perkins, A., Nelson, K. J., Parsonage, D., Poole, L. B. & Karplus, P. A. Peroxiredoxins: guardians against oxidative stress and modulators of peroxide signaling. Trends in Biochemical Sciences 40, 435–445 (2015).

51. Rulísek, L. & Vondrásek, J. Coordination geometries of selected transition metal ions (Co2+, Ni2+, Cu2+, Zn2+, Cd2+, and Hg2+) in metalloproteins. J Inorg Biochem 71, 115–127 (1998).

52. Li, X., Eyles, S. J. & Thompson, L. K. Hydrogen exchange of chemoreceptors in functional complexes suggests protein stabilization mediates long-range allosteric coupling. J Biol Chem 294, 16062–16079 (2019).

53. Reyes, G. I., Flack, C. E. & Parkinson, J. S. The structural logic of dynamic signaling in the Escherichia coli serine chemoreceptor. Protein Sci 33, e5209 (2024).

54. Tama, F. & Sanejouand, Y.-H. Conformational change of proteins arising from normal mode calculations. Protein Engineering 14, 1–6 (2001).

55. Gerstein, M., Lesk, A. M. & Chothia, C. Structural Mechanisms for Domain Movements in Proteins. Biochemistry 33, 6739–6749 (1994).

56. Ortega, D. R. et al. A phenylalanine rotameric switch for signal-state control in bacterial chemoreceptors. Nat Commun 4, 2881 (2013).

57. Burt, A. et al. Complete structure of the chemosensory array core signalling unit in an E. coli minicell strain. Nat Commun 11, 743 (2020).

58. Sies, H., Mailloux, R. J. & Jakob, U. Fundamentals of redox regulation in biology. Nat Rev Mol Cell Biol 25, 701–719 (2024).

59. Koshland, D. E. The structural basis of negative cooperativity: receptors and enzymes. Curr Opin Struct Biol 6, 757–761 (1996).

60. Cooperative Signaling among Bacterial Chemoreceptors | Biochemistry. https://pubs.acs.org/doi/10.1021/bi050567y.

61. Cho, Y. R. et al. Cooperativity and anti-cooperativity between ligand binding and the dimerization of ristocetin A: asymmetry of a homodimer complex and implications for signal transduction. Chemistry & Biology 3, 207–215 (1996).

62. Bartelli, N. L. & Hazelbauer, G. L. Differential backbone dynamics of companion helices in the extended helical coiled-coil domain of a bacterial chemoreceptor. Protein Sci 24, 1764– 1776 (2015).

63. Akkaladevi, N., Bunyak, F., Stalla, D., White, T. A. & Hazelbauer, G. L. Flexible Hinges in Bacterial Chemoreceptors. Journal of Bacteriology 200, 10.1128/jb.00593-17 (2018).

64. Elliott, L. G., Simpkin, A. J. & Rigden, D. J. ABCFold: easier running and comparison of AlphaFold 3, Boltz-1, and Chai-1. Bioinformatics Advances 5, vbaf153 (2025).

65. Simpkin, A. J. et al. Slice’N’Dice: maximizing the value of predicted models for structural biologists. Acta Crystallogr D Struct Biol 81, 105–121 (2025).

66. Wohlwend, J. et al. Boltz-1 Democratizing Biomolecular Interaction Modeling. 2024.11.19.624167 Preprint at 10.1101/2024.11.19.624167 (2025).

67. STARANISO anisotropy & Bayesian estimation server. https://staraniso.globalphasing.org/cgi-bin/staraniso.cgi.

68. Kulczyk, A. W. Artificial intelligence and the analysis of cryo-EM data provide structural insight into the molecular mechanisms underlying LN-lamininopathies. Sci Rep 13, 17825 (2023).

69. Kulczyk, A. W., McKee, K. K. & Yurchenco, P. D. Cryo-EM reveals molecular mechanisms underlying the inhibitory effect of netrin-4 on laminin matrix formation. Nat Commun 16, 7256 (2025).

70. Kulczyk, A. W. et al. Cryo-EM reveals the molecular basis of laminin polymerization and LN-lamininopathies. Nat Commun 14, 317 (2023).

71. Kulczyk, A. W. et al. Cryo-EM structure of Shiga toxin 2 in complex with the native ribosomal P-stalk reveals residues involved in the binding interaction. J Biol Chem 299, 102795 (2023).

72. Kulczyk, A. W., Moeller, A., Meyer, P., Sliz, P. & Richardson, C. C. Cryo-EM structure of the replisome reveals multiple interactions coordinating DNA synthesis. Proceedings of the National Academy of Sciences 114, E1848–E1856 (2017).

73. Punjani, A., Rubinstein, J. L., Fleet, D. J. & Brubaker, M. A. cryoSPARC: algorithms for rapid unsupervised cryo-EM structure determination. Nat Methods 14, 290–296 (2017).

74. DiIorio, M. C. & Kulczyk, A. W. A Robust Single-Particle Cryo-Electron Microscopy (cryo-EM) Processing Workflow with cryoSPARC, RELION, and Scipion. J Vis Exp 10.3791/63387 (2022) doi:10.3791/63387.

75. Adams, P. D. et al. PHENIX: a comprehensive Python-based system for macromolecular structure solution. Acta Crystallogr D Biol Crystallogr 66, 213–221 (2010).

